# Aging increases ovarian cancer growth, metastasis, and immunosuppression that can be alleviated by inhibiting hedgehog signaling

**DOI:** 10.64898/2025.12.23.695206

**Authors:** Asha Kumari, Mohamed Halaby Elbahoty, Resha Rajkarnikar, Khushi Sureja, Manan Virendra Nayyar, Mehri Monavarian, Liz Macias Quintero, Katherine Fuh, Daniel J Tyrrell, Lalita A Shevde, Sudarshan Anand, Camilla Margaroli, Karthikeyan Mythreye

## Abstract

Ovarian cancer incidence and mortality increase with age, yet how aging shapes tumor progression and the immune microenvironment remains poorly defined. Using orthotopic syngeneic models of distinct cellular origins (ovarian surface epithelial and fallopian tube-derived) in young versus aged mice, we show that aged hosts exhibit higher tumor burden, metastasis and ascites. Follicle depletion in young mice did not recapitulate these effects, indicating contributions beyond hormonal decline. Spatial transcriptomics revealed distinct age dependent intratumoral heterogeneity, with Hedgehog signaling enrichment in CD45^+^ cells from aged tumors, alongside elevated CD206^+^ tumor-associated macrophages and FoxP3^+^ regulatory T cells. Pharmacologic Hedgehog inhibition in aged mice suppressed tumor growth, reduced metastasis, and decreased CD206^+^ macrophages and FoxP3^+^ T cells while preserving CD8^+^T cells. In human ovarian cancer, Hedgehog activation correlated with immunosuppressive and immune checkpoint resistance signatures. We propose Hedgehog inhibition as an immunomodulatory strategy for Hedgehog activated or post menopausal ovarian cancer.

## Introduction

Age represents one of the strongest risk factors across most cancer types, with both cancer incidence and mortality rates increasing dramatically in patients over 65 years. However, the mechanisms by which physiological aging contributes to tumor progression remain incompletely understood. Aging is associated with broad alterations in the tissue microenvironment that include but are not limited to changes in extracellular matrix composition, accumulation of senescent cells, and shifts in immune cell populations, all of which may create a pro-tumorigenic environment. Ovarian cancer represents a particularly compelling model, as incidence and mortality increase sharply after menopause, with a 26% higher mortality among aged patients compared to other cancer types^1,2^. In this context, hormonal decline has historically been considered a driver of increased susceptibility, yet whether broader age-associated microenvironmental changes contribute independently remains unclear. High-grade serous ovarian carcinoma (HGSOC), the most lethal gynecologic malignancy, often originates in the fallopian tube fimbriae as serous tubal intraepithelial carcinoma (STIC) lesions, which shed cells that engraft on the ovarian surface and form the bulk primary tumor before peritoneal dissemination^3–5^. Other ovarian cancer subtypes including endometrioid, mucinous, and clear cell, typically arise directly from ovarian tissue or other pelvic structures. Regardless of cell of origin, the ovary serves as an obligate site of bulk disease establishment for HGSOC, which is the rationale for risk-reducing salpingo-oophorectomy in BRCA1/2 carriers, a procedure that substantially reduces ovarian cancer incidence and all-cause mortality^6–9^. This makes the aging ovarian microenvironment a potential key determinant of disease progression. Despite this clinical reality, most preclinical ovarian cancer studies employ young animal mouse models, that bypass ovarian age-specific tumor-host interactions, limiting our understanding of how physiological aging influences tumor behavior.

Older ovarian cancer patients also experience poorer outcomes, suggesting that age-associated changes in the ovarian and peritoneal cavity may contribute to tumor progression and treatment response to therapies ^10^. Ovarian aging also extends beyond follicle decline, including extracellular matrix composition changes ^11^ accumulation of senescent and multinucleated ovarian stromal cells and macrophages ^12^ and significant shifts in immune cell populations. These immune shifts in mouse models, include higher levels of monocyte recruitment followed by increased alternatively activated macrophage (M2) populations ^13^ and a shift towards adaptive immunity ^14^. These changes in the immune milieu may predispose aged ovaries to increased tumor burden. Whether hormonal decline alone accounts for age-associated tumor progression, or whether broader aging-related changes are required, remains unclear. Prior work in 85 week old female mouse using intraperitoneal injections, showed increased tumor burden and tumor infiltrating lymphocytes (TIL) with altered B cell related pathways in the peritoneal adipose tissues^15^. However, how tumor growth, metastasis, and the immune landscape differ in aged versus young ovarian microenvironments has not been systematically defined. Similarly, survival analysis in a genetically engineered mouse model indicates that nulliparous aged mice had shorter survival compared to young and multiparous mice^16^ indicating that parity, and the likely associated hormonal changes influence tumor outcomes.

Here, we compared tumor progression and immune composition differences and outcomes in 60-65 week-old female mice, which represent reproductive decline corresponding to menopause in women (∼51 years) ^17–19^, versus young mice (controlling for parity) and follicle depleted mice. We find that menopausal hosts support tumor progression significantly more than younger hosts. Using spatial transcriptomics, we analyzed age-associated differences in the tumor immune microenvironment and identified Hedgehog signaling as a pathway enriched in immune cells from aged tumors, which showed a concurrent increase in a spectrum of immunosuppressive cell populations. Notably, pharmacologic inhibition of Hedgehog signaling reduced tumor progression and the presence of these same immunosuppressive populations. Finally, in human ovarian cancer datasets, Hedgehog pathway activation correlated with an immunosuppressive microenvironment, including higher myeloid suppressor markers, lower cytotoxic effector signatures, and immune checkpoint blockade resistance programs. Together, these findings demonstrate that physiological aging, rather than follicular depletion alone, is associated with accelerated tumor growth and an immunosuppressive microenvironment, and implicate the Hedgehog signaling pathway as a potential target for immune remodeling in post-menopausal hosts.

## Results

### Age-associated changes, rather than follicular depletion alone, promote ovarian cancer progression

To investigate how physiological ovarian aging affects tumor growth and metastasis in ovarian and fallopian tube-derived cancer, we compared tumor growth and metastasis in young (6-8 weeks) versus aged (60-65 weeks) nulliparous C57BL/6 mice to eliminate physiological and hormonal changes associated with pregnancy ^20^. Here, ‘young’ and ‘aged’ refer exclusively to the chronological age of the host at the time of tumor implantation, not to tumor stage or duration of disease. Aged mice (60-65 weeks, ∼15 months) correspond approximately to peri/post-menopausal women, while young mice (6-8 weeks) correspond to reproductively active pre-menopausal women^19,20^. Ovaries from aged host mice showed predicted changes, including a 10.5-fold reduction in follicle count and decreased expression of ovarian function markers (1.6-fold lower *Inha* and 3.4-fold lower *Amh* mRNA) compared to ovaries from young mice (Supplementary Figure 1A, i-iii). To quantify differences in tumor growth in the hosts we orthotopically implanted two complementary and distinct syngeneic tumor cell models representing the two proposed cellular origins of epithelial ovarian cancer: ID8*Trp53^−/−^*(ovarian surface epithelial-derived) and PPNM (fallopian tube-derived; *p53^R172H^ Pten^−/−^Nf1^−/−^Myc^OE^*) into the mice ovarian bursa. The PPNM mutational profile recapitulates alterations recurrent in HGSOC patients ^21^, including TP53 mutation (>95%), MYC amplification (∼20-30%), and PTEN or NF1 loss (∼5-8% each) ^22^. Tumor growth kinetics were monitored by bioluminescence imaging in ID8*Trp53^−/−^*implanted mice (Supplementary Figure 1B, i–ii) and by palpation in PPNM-implanted mice. BLI signal was comparable between young and aged hosts at early timepoints (Days 7–18), with divergence beginning around Day 18–21 (Supplementary Figure 1B, i-ii), indicating that the age-dependent phenotype reflects differential tumor progression in the host environment rather than differences in initial tumor cell establishment in the bursa. Because mice were euthanized at a uniform endpoint, we quantified progression using a time-to-threshold analysis. The event was defined as the first attainment of ≥2.77 cm² peritoneal spread and yielded Kaplan Meier style estimates of the probability of remaining below threshold over time. Aged hosts reached this threshold significantly earlier, with a 3.7-fold higher hazard of progression. (Supplementary Figure 1B, iii). At endpoint (day 55 for ID8*Trp53^−/−^*, day 42 for PPNM), aged mice displayed significantly greater tumor burden across all metastatic sites (Supplementary Figure 1B, v–vi). In the ID8*Trp53^−/−^*model, aged hosts showed increased ovarian and omental tumor mass (1.8-fold and 1.2-fold respectively), ascites volume (1.6-fold), and total peritoneal/mesenteric burden (1.8-fold) relative to young hosts (Figure 1A; Supplementary Figure 1C, i–ii). In the PPNM model, age-related differences were even more pronounced, with 2.3-fold greater ovarian tumor mass, 6.2-fold greater omental tumor mass, 9.1-fold higher ascites volume, and 2.7-fold higher total tumor burden in aged compared to young mice (Figure 1D; Supplementary Figure 1C, iii–iv). Histological analysis revealed that tumors from aged hosts had higher proliferation rates as determined by PCNA staining. Specifically, ovarian tumors were 1.6-fold more proliferative in both ID8*Trp53^−/−^*and PPNM, and omental tumors were 1.4-fold for ID8*Trp53^−/−^* and 1.2-fold for PPNM as compared to young hosts (Figure 1B-C, E-F).

**Figure 1.**
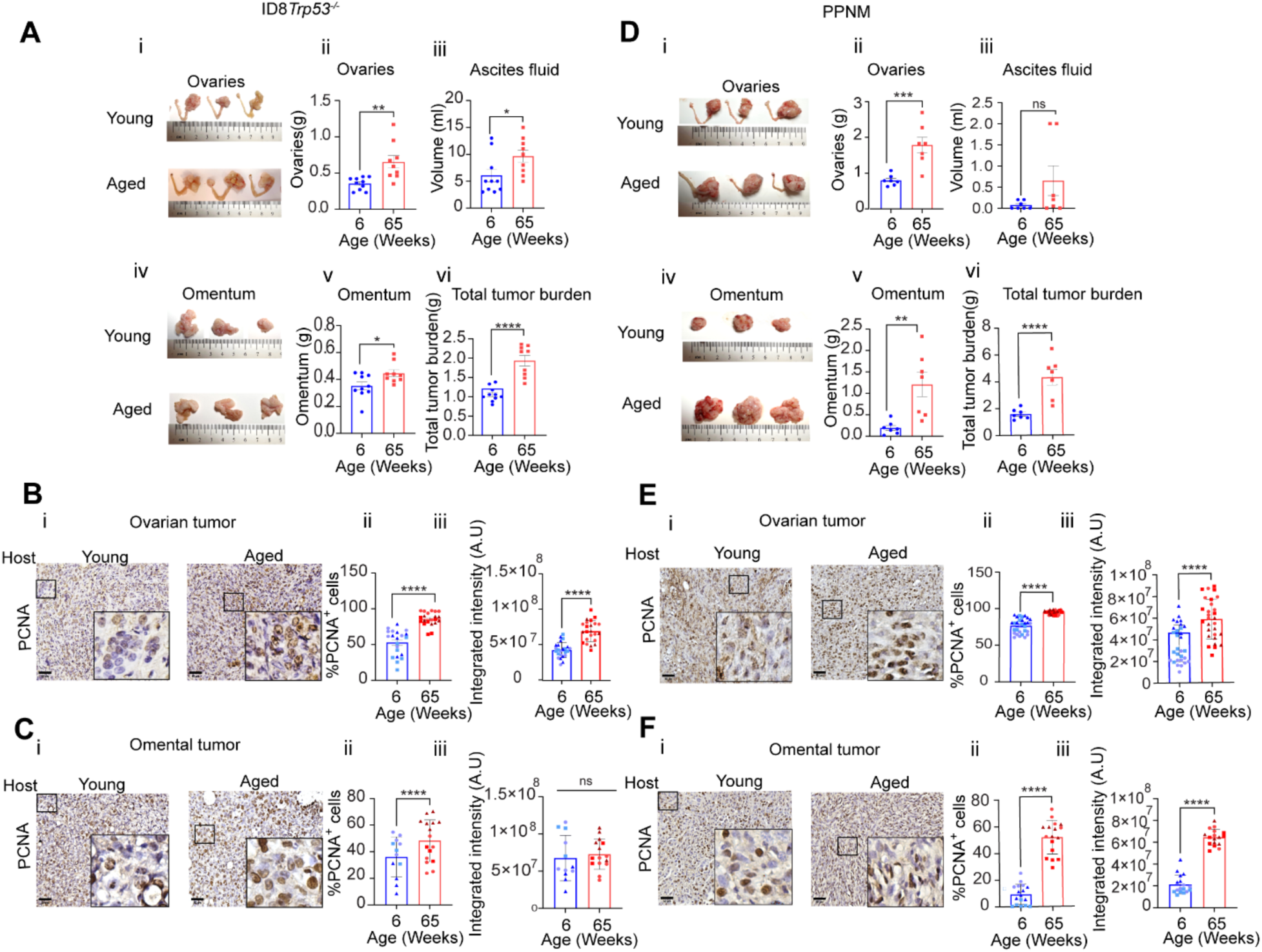
**(A)** (i)&(iv) Representative images of tumor burden in the ovaries and omentum of mice of the indicated age groups. (ii) Quantification of ovary weights (iii) ascites fluid volume (v) omental weights (vi) total tumor burden, on day 55 post intrabursal implantation of ID8*Trp53^−/−^*cells in young and aged mice ovaries (n=9). **(B-C)** Immunohistochemical analysis of proliferating nuclear antigen (PCNA) in **(B)**(i) ovarian and **(C)**(i) omental tumors from young or aged mice (ID8*Trp53^−/−^*) (ii)&(iii) Percentage and integrated intensity of PCNA-positive cells in ovarian and omental tumors (n=3). **(D)**(i)&(iv) Representative images of tumor burden in the ovaries and omentum of mice of the indicated age groups. (ii) Quantification of ovary weights (iii) ascites fluid volume (v) omental weights (vi) total tumor burden on day 42 following intrabursal implantation of murine fallopian tube-derived PPNM cells in young and aged mice ovaries (n=8). (**E-F)** PCNA immunohistochemistry of **(E)**(i) ovarian and **(F)**(i) omental tumors from young or aged mice (PPNM) (ii-iii) percentage and intensity of PCNA-positive cells in ovarian and omental tumors (n=3). Color codes and symbols in the graphs represent the number of images analyzed per mouse. All data are presented as mean±SEM, *p<0.05,**p<0.01, ***p<0.001, ****p<0.0001, unpaired t test.

To determine whether follicular depletion and associated hormonal changes alone could account for increased tumor growth, we treated young mice with 4-vinylcyclohexene diepoxide (VCD), which selectively depletes ovarian follicles without other age-associated changes. VCD treatment successfully reduced follicle number by 3.7-fold and decreased *AMH* and *INHA* expression by 14-fold and 10-fold, respectively (Supplementary Figure 1D, i-iii). ID8*Trp53^−/−^* cells implanted into the ovarian bursa of follicle depleted ovaries (VCD pretreated) and follicle replete ovaries (vehicle-treated) showed no significant difference at endpoint (day 67 post-implantation) (Supplementary Figure 1D, iv). We confirmed no significant differences in ovarian or omental tumor weights, ascites volume, or total tumor burden between groups (Supplementary Figure 1D, v-xi). These findings demonstrate that follicle depletion in the ovaries alone does not support tumor growth; rather, physiological age in mice strongly accelerates tumor growth in the ovaries, and metastasis to the omentum and peritoneal cavity, thereby reducing overall survival. To further test whether the increased tumor burden in aged mice is preserved when tumor cells encounter the peritoneal compartment directly, the dominant site of advanced HGSOC disease spread, PPNM cells were injected intraperitoneally (i.p) into young (6-8 weeks) and aged (60-65 weeks) mice. PPNM cells were not found to grow significantly differently in the ovary when implanted through the peritoneum (Supplementary Figure 1E, iii). However aged mice showed higher total intraperitoneal tumor burden, omental tumor weight, and ascites volume compared to young mice (Supplementary Figure 1E, v-viii).Together, the reproducibility of the aged tumor burden phenotype across two tumor models (ovarian surface epithelial-derived ID8*Trp53^−/−^* and fallopian tube-derived PPNM) and two delivery routes (intrabursal and intraperitoneal), is more consistent with a systemic host aging effect than a compartment or delivery route restricted one.

### Spatial transcriptomics reveal distinct intratumoral heterogeneity patterns in ovarian tumors from young versus aged hosts

To understand the spatial organization of the tumor microenvironment in young and aged hosts, and define intra and intertumoral differences, we performed spatial transcriptomics using digital spatial profiling (DSP). Regions from ovarian tumors from young and aged mice were segmented into tumor cells (CK8^+^), immune cells (CD45^+^), and endothelial cells (CD31^+^) by multiplex immunofluorescence (Figure 2 A-B). Given the well-documented immune alterations associated with ovarian cancer and ovarian aging, and the critical role of immune cells in ovarian cancer tumor progression, we focused our analysis for this study on comparing CD45^+^ enriched regions versus CD45^+^ depleted regions (n=6 ROIs from each category). Within young host tumors, CD45^+^ depleted regions showed enrichment in hallmark pathways associated with metabolic reprogramming, including adipogenesis, oxidative phosphorylation, and fatty acid metabolism (Figure 2C, i). Top upregulated genes in these regions included *Fxyd3* (FXYD Domain Containing Ion Transport Regulator 3), *Prrl5l* (Proline Rich 15 Like), and *Flrt1* (Fibronectin Leucine Rich Transmembrane Protein 1) (Supplementary Table A). KEGG pathway analysis revealed enrichment in glycan biosynthesis pathways (Supplementary Figure 2A, i). In contrast, CD45^+^ enriched regions in the same young host, tumors displayed allograft rejection, epithelial-mesenchymal transition (EMT), E2F targets, and inflammatory response hallmark pathways (Figure 2C, i). Top upregulated genes included *Thbs1* (thrombospondin-1), *Col8a1* (collagen type VIII alpha-1 chain), and *Adgre1* (Adhesion G Protein-Coupled Receptor E1) (Supplementary Table A). KEGG pathways related to tuberculosis, malaria, and immune deficiency were enriched in these regions (Supplementary Figure 2A, ii).

**Figure 2.**
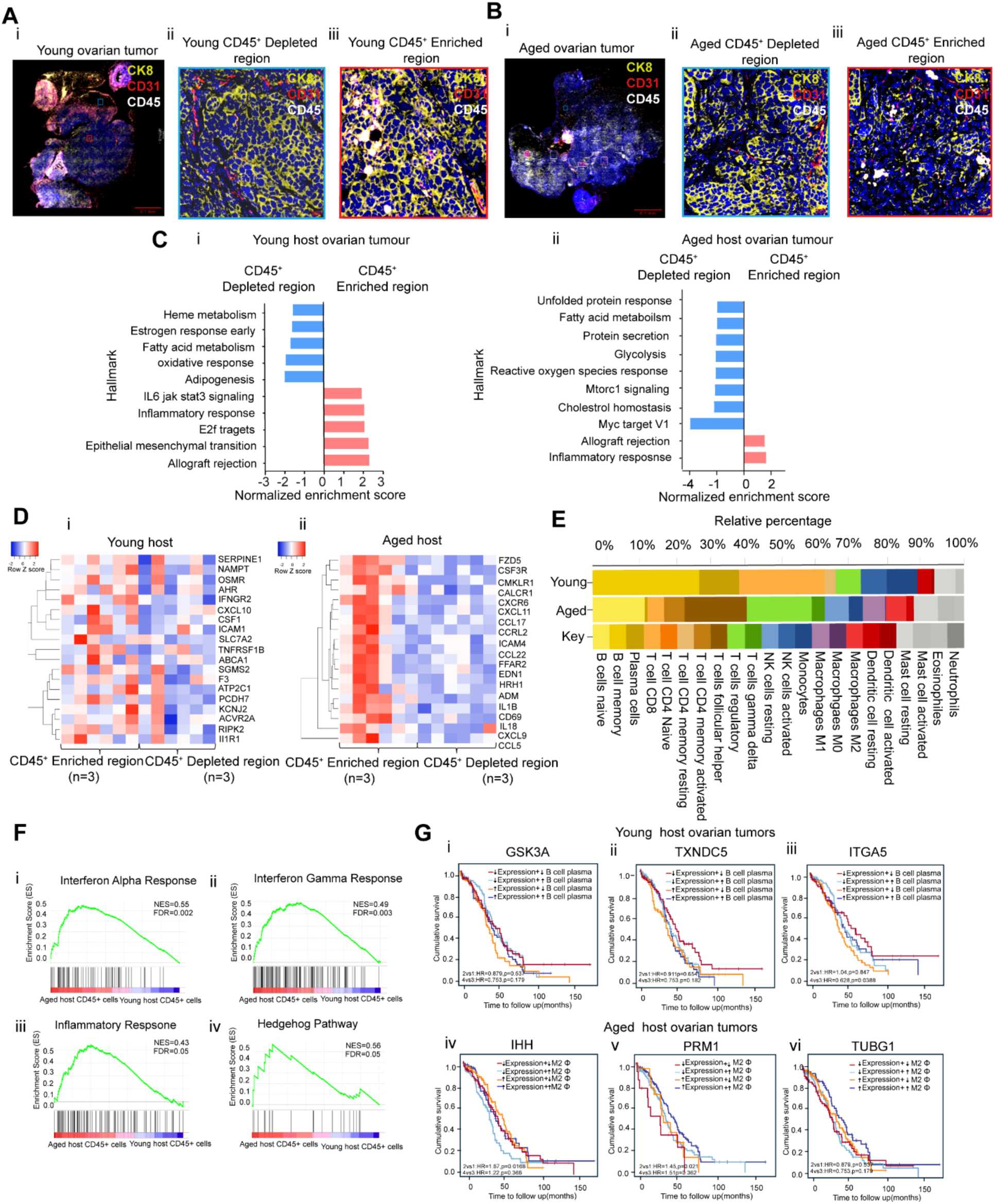
**(A-B)** Representative multiplex immunofluorescence images of an ovarian tumor from young or aged hosts. Tumor cells are identified as CK8⁺(yellow), immune cells as CD45⁺ (white), endothelial cells as CD31⁺(red), and nuclei (blue). Magnified panels highlight representative regions of interest (ROIs) selected from CD45⁺depleted (<10–15 CD45^+^ cells determined by) or CD45⁺enriched areas (> 50 CD45^+^ cells) for Spatial GeoMx analysis (n=3). **(C)** Intra-tumoral Gene Set Enrichment Analysis (GSEA) comparing CD45⁺depleted or CD45⁺-enriched regions from (i) young and (ii) aged hosts tumors using normalized enrichment scores (NES) against MSigDB Hallmark gene sets (FDR<0.05). **(D)** Heatmaps depicting unique inflammatory genes contributing to the ‘Inflammatory Hallmark’ enrichment in ovarian tumors from (i) young (ii) aged hosts. (**E)** CD45^+^ cells deconvolution using CIBERSORTX from young and aged host tumors. **(F)**(i-iv) GSEA of CD45⁺-enriched regions compared between young and aged hosts using Hallmark gene sets (FDR<0.05). **(G)** Kaplan-Meier survival analysis of HGSOC patients from TIMER 2.0, stratified by median expression of each gene (high = above median; low = below median; standard TCGA approach). (i-iii) Genes upregulated in young host ovarian tumors (*GSK3A, TXNDC5, ITGA5*); high expression in patients is associated with higher B cell infiltration and longer survival. (iv-vi) Genes upregulated in aged host ovarian tumors (*IHH, PRM1, TUBG1*); high expression in patients is associated with higher M2 macrophage infiltration and shorter survival.

In contrast to tumors from young hosts, ovarian tumors from aged hosts displayed a distinct pattern of intratumoral heterogeneity. CD45^+^ cells depleted regions showed enrichment for MYC targets V1, cholesterol homeostasis, and mTORC1 signaling hallmark pathways (Figure 2C, ii). Top upregulated genes included *Clu* (Clusterin), *Gpc1* (Glypican-1), and *TgfbI* (Transforming Growth Factor Beta Induced) (Supplementary Table B). KEGG pathways related to protein processing were enriched in these regions (Supplementary Figure 2A, iii). CD45^+^ cells enriched regions in tumors from aged hosts were characterized by inflammatory response, allograft rejection, and KRAS signaling hallmark pathways (Figure 2C, ii). Top upregulated genes included *Hmgcs2* (3-Hydroxymethylglutaryl-CoA Synthase 2), and *Lyz2* (Lysozyme 2) (Supplementary Table B). KEGG pathway analysis showed enrichment in chemokine signaling, cytokine-cytokine receptor interaction, leukocyte transendothelial migration, and natural killer cell cytotoxicity pathways (Supplementary Figure 2A, iv).

Comparison of CD45^+^ depleted regions between the young and old ovarian tumor groups revealed three common hallmark pathways: fatty acid metabolism, androgen response, and heme metabolism (Supplementary Figure 2A, v) implicating these pathways as host age independent. CD45^+^ enriched regions from the young and old host ovarian groups shared the inflammatory hallmark (Supplementary Figure 2A, vi). However, even in the shared inflammatory signature, the genes driving this hallmark were non-overlapping, with 19 unique genes in each age group (Figure 2D, i-ii). For example, tumors from young hosts expressed *Cxcl10*, associated with anti-tumor immunity, whereas aged tumors expressed chemokines linked to immunosuppression, including *Ccl5*, *Ccl22*, and *Ccl17*. These findings indicate that while tumors from both young and aged hosts exhibit intratumoral heterogeneity, the underlying molecular programs, particularly those governing inflammation, are fundamentally distinct, suggesting potential age-associated differences in immune cell composition and function.

### Immune cells from tumors of aged hosts are immunosuppressive with Hedgehog pathway signatures

To determine whether these pathway differences reflect altered immune cell composition, we performed deconvolution using CIBERSORTX^23^.Young host tumors contained higher proportions of memory B cells and CD8^+^ T cells. In contrast, aged host tumors were dominated by regulatory T cells (18% vs 4% in young) and CD4^+^ follicular T helper cells (Figure 2E). Notably, tumor associated M2 macrophages (TAM), that are known drivers of immunosuppression in ovarian cancer ^24,25^ were markedly elevated in tumors from aged hosts (5% vs 1% in young). To investigate the functional programs active in these immune populations, we performed GSEA on CD45^+^ cells themselves. In aged host tumors CD45^+^ cells were characterized by IFN-α, IFN-γ, inflammatory response, and Hedgehog signaling pathways (Figure 2F, i-iv). In contrast, CD45^+^ cells from young host tumors showed enrichment in EMT, Notch signaling, and estrogen response early, though these did not reach statistical significance. Consistent with these compositional differences, the top upregulated genes in CD45^+^ cells from tumors from young hosts include *Gsk3a*, *Txndc5*, and *Itga5*; genes that are associated with B cell infiltration and prolonged survival in ovarian cancer patients (Supplementary Figure 2B, ii Figure 2G, i-iii). Notably, Jchain (immunoglobulin joining chain, essential for polymeric IgA and IgM secretion) was also among the genes upregulated in young host tumors by differential expression analysis (Supplementary Figure 2B, i), consistent with an active humoral immune response in this group. Conversely, top upregulated genes in CD45^+^ cells from aged tumors include *Prm1*, *Ihh* (Indian Hedgehog), and *Tubg1*, are associated with M2 macrophage infiltration as shown in other models previously ^26,27^ and poor survival outcomes in ovarian cancer patients (Supplementary Figure 2B, ii, Figure 2G, iv-vi). Together, these analyses reveal that aged host tumors harbor immunosuppressive cell populations with Hedgehog ligand and pathways, identifying this as a potential modifiable target.

### Tumors from aged hosts are enriched for immunosuppressive macrophages and regulatory T cells

To phenotypically validate CIBERSORTX predictions, we quantified immunosuppressive cell populations in ovarian and omental tumors, the latter being a preferred metastatic site in ovarian cancer ^28^. We first assessed CD45^+^ cells co-expressing Arginase 1 (Arg1), the L-arginine–metabolizing enzyme whose depletion in the tumor microenvironment impairs T cell and NK cell proliferation and cytotoxic function^29,30^. In aged hosts, ovarian tumors showed 2.8-fold higher CD45^+^Arg1^+^ cells in the ID8*Trp53^−/−^* model (Figure 3A) and 2.42-fold higher in the PPNM model (Figure 3B) compared to ovarian tumors in young hosts. Similarly, omental tumors from aged hosts showed 3.52-fold (ID8*Trp53^−/−^*, Figure 3I) and 2.08-fold (PPNM, Figure 3J) higher CD45^+^Arg1^+^ cells. Total CD45^+^ infiltration followed similar trends (Supplementary Figure 3A, i-iv). Since Arg1^+^ expressing monocytes/macrophages (CD68^+^) promote tumor progression and immune evasion in ovarian and other cancers^31,32^, we specifically quantified CD68^+^Arg1^+^ cells. Ovarian tumors in aged hosts showed 3.7-fold and 2-fold higher CD68^+^Arg1^+^ cells compared to young hosts in ID8*Trp53^−/−^* and PPNM models, respectively (Figure 3C,D). CD68^+^Arg1^+^ cells were also numerically higher in aged omental tumors, reaching statistical significance in the PPNM model (Figure 3L). The balance between CD80^+^ and CD206^+^ macrophages is crucial in determining the local immune milieu and prognosis in ovarian cancer^33,34^. Aged host tumors showed 2.69-fold higher CD206^+^ cells in ovarian tumors (ID8*Trp53^−/−^*, Figure 3E) and 2.26-fold higher in the PPNM model (Figure 3F). Omental tumors showed similar elevations in CD206^+^ cells in aged hosts (Figures 3M, 3N). Total CD68^+^ macrophages were also significantly elevated in aged ovarian tumors and trended higher in omental tumors (Supplementary Figure 3A, v-viii), consistent with both increased macrophage recruitment and preferential CD206^+^ M2 expansion in the aged TME. Further supporting these findings, ascites fluid from aged mice (ID8*Trp53*^−/−^) showed a 2.9-fold higher ratio of F4/80^+^CD206^+^ to F4/80^+^CD80^+^ macrophages (Supplementary Figure 3B, iv, v).

**Figure 3.**
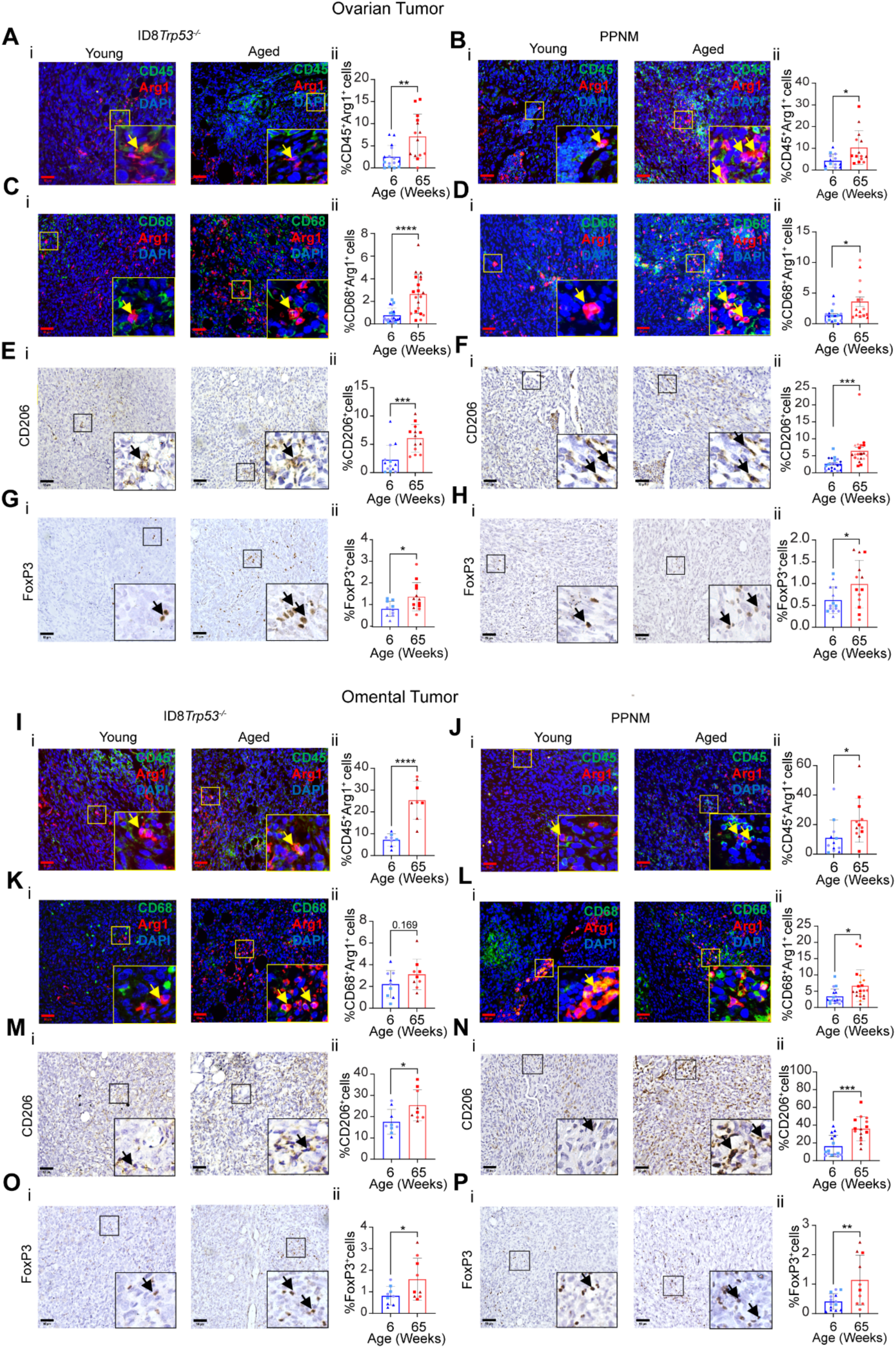
**(A-B)** Representative immunofluorescence images from indicated ovarian tumors showing CD45^+^(green), Arginase-1 (Arg1^+^) (red), and DAPI (blue). Adjacent graphs show quantification of percentage CD45^+^Arg1^+^ cells in ovarian tumors from indicated models. **(C-D)** Representative immunofluorescence images from tumors from the ovary showing CD68^+^(green), Arg1^+^(red), and DAPI (blue) cells. Adjacent graphs show quantification of percentage CD68^+^Arg1^+^ cells from both models (ID8*Trp53^−/−^* and PPNM) **(E-F)** Representative CD206^+^ immunohistochemistry of ovarian tumors with adjacent quantification of percentage CD206^+^cells in indicated tumors from both models. **(G-H)** Representative FoxP3^+^ immunohistochemistry images in ovarian tumors with adjacent quantification of percentage Foxp3^+^ cells in both models. **(I-J)** Representative immunofluorescence images from omental tumors showing CD45^+^(green), Arg1^+^(red), and DAPI (blue). Adjacent graphs show quantification of percentage CD45^+^Arg1^+^ cells in omental tumors. **(K-L)** Representative immunofluorescence images from omental tumors showing CD68^+^(green), Arg1^+^(red), and DAPI (blue) cells with adjacent graphs showing quantification of percentage CD68^+^Arg1^+^ cells in the omental tumors. **(M-N)** Representative CD206^+^ immunohistochemistry of omental tumors with adjacent quantification of percentage CD206^+^cells in the tumors. **(O-P)** Representative FoxP3^+^ immunohistochemistry images in omental tumors with adjacent quantification of percentage Foxp3^+^ cells in both models. Color codes and symbols indicate the number of images analyzed per mouse in each group. All data are Mean±SEM (n=3), *p<0.05, **p<0.01, *** p<0.001, ****p<0.0001, unpaired t test.

We next evaluated regulatory T cells, a critical population known to inhibit effective anti-tumor immunity ^35^ and predicted to be elevated from our deconvolution analysis (Figure 3H). FoxP3^+^ cells were 3 to 3.4-fold higher in aged host ovarian tumors across both cell models (Figures 3G, 3H and Supplementary Figure 3A xiii-xvi) and 2.28-fold higher in omental tumors (Figures 3O, 3P and Supplementary Figure 3A xxii-xxv). Consistent with tumor tissue findings, ascites fluid from aged mice showed 2.5-fold higher CD25^+^FoxP3^+^ cells (Supplementary Figure 3B, ix, x). CD8^+^ cytotoxic T cells were significantly reduced in aged host ovarian and omental tumors from both ID8*Trp53*^−/−^ and PPNM models (Supplementary Figure 3A, ix-xii) independent of whether cells were delivered intrabursally or intraperitoneally (Supplementary Figure 3C, i-ii). Omental tumors from PPNM cells delivered intraperitoneally also showed a similar and significant trend of elevated CD206^+^M2 macrophages and Tregs (Supplementary Figure 3C) indicating a shift toward immunosuppression in tumors from aged hosts regardless of route of delivery of tumor cells.

To further characterize the myeloid immunosuppressive compartment, we analyzed ascites flow cytometry data to quantify monocytic MDSCs (M-MDSCs: CD11b^+^Ly6G^-^Ly6C^high^) and polymorphonuclear MDSCs (PMN-MDSCs: CD11b^+^Ly6G^+^Ly6C^low^), along with double-positive:CD11b^+^Ly6G^+^Ly6C^+^ and CD11b^+^Ly6G^-^Ly6C^-^. We found that M-MDSCs:CD11b^+^Ly6G^-^Ly6C^high^and double-positive CD11b^+^Ly6G^+^Ly6C^+^ populations trended towards being higher in aged mice, though differences did not reach statistical significance (Supplementary Figure 3B, xi-xv); however, CD11b^+^Ly6G^-^Ly6C^-^ cells were significantly higher in aged hosts as compared to younger hosts (Supplementary Figure 3B, xii). Together, these findings demonstrate that the aged tumor microenvironment is dominated by immunosuppressive cell populations including M2 macrophages and regulatory T cells, across primary (ovary) and metastatic (omentum) sites, providing a cellular basis for the enhanced tumor growth observed in aged hosts.

### Hedgehog inhibition suppresses ovarian tumor growth and metastasis in aged models

Based on the immune suppressive cell populations, accelerated tumor growth and predominance of Hedgehog signaling in immune cells from aged host tumors, we aimed to test the therapeutic benefit of inhibiting Hedgehog (HH) signaling pathway in aged hosts using Vismodegib (Figures 4A,4B). Mice were euthanized at a uniform endpoint, and thus disease progression was assessed using a time-to-threshold probability analysis of disease progression using abdominal girth changes. This yielded Kaplan-Meier-style estimates of the probability of remaining below the abdominal girth threshold. Vismodegib-treated mice showed delayed disease progression, remaining below threshold until day 41 compared to day 16 in vehicle-treated mice (Figure 4C). Vismodegib treatment also reduced overall metastatic burden (Figure 4D-F) as determined by the number of mice that developed measurable ascites (33.3% of vismodegib vs 88.8% of vehicle-treated mice, Figure 4D), diaphragmatic metastases (66.6 % of vismodegib vs 87.5% of vehicle-treated mice Figure 4E), and peritoneal wall metastases (22.2% of vismodegib mice vs 50% of vehicle-treated mice Figure 4F). Ovarian and omental tumor weights trended lower in vismodegib treated mice, ovarian weights did not reach statistical significance (Figure 4G-H). Tumor proliferation was markedly reduced as PCNA-positive cells were 1.47-fold lower in ovarian tumors and 1.86-fold lower in omental tumors from vismodegib treated mice, with corresponding reductions in staining intensity (1.5-fold and 2.15-fold, respectively, Figure 4G,4H). Vismodegib was well-tolerated, with no significant differences in body weight or clinical signs (activity, grooming, hunched posture, piloerection, diarrhea) between vismodegib and vehicle-treated mice (Supplementary Figure 4A). Systemic Hedgehog pathway inhibition was also confirmed by qRT-PCR in mesentery tissue, which showed downregulation of Hedgehog target genes inn vismodegib treated mice relative to vehicle-treated controls (Supplementary Figure 4B). To determine whether the in vivo anti-proliferative effects of vismodegib reflect direct tumor cell cytotoxicity, we treated ID8*Trp53^−/−^* cells in vitro across a range of vismodegib concentrations. Minimal sensitive to vismodegib, with IC50 values>49 µM (Supplementary Figure 4C), was observed. These findings demonstrate that HH signaling inhibition delays tumor progression and reduces metastatic spread in aged hosts and the reduced tumor proliferation observed in vivo (Figure 4G,H) is likely not driven by direct tumor cell cytotoxicity.

**Figure 4.**
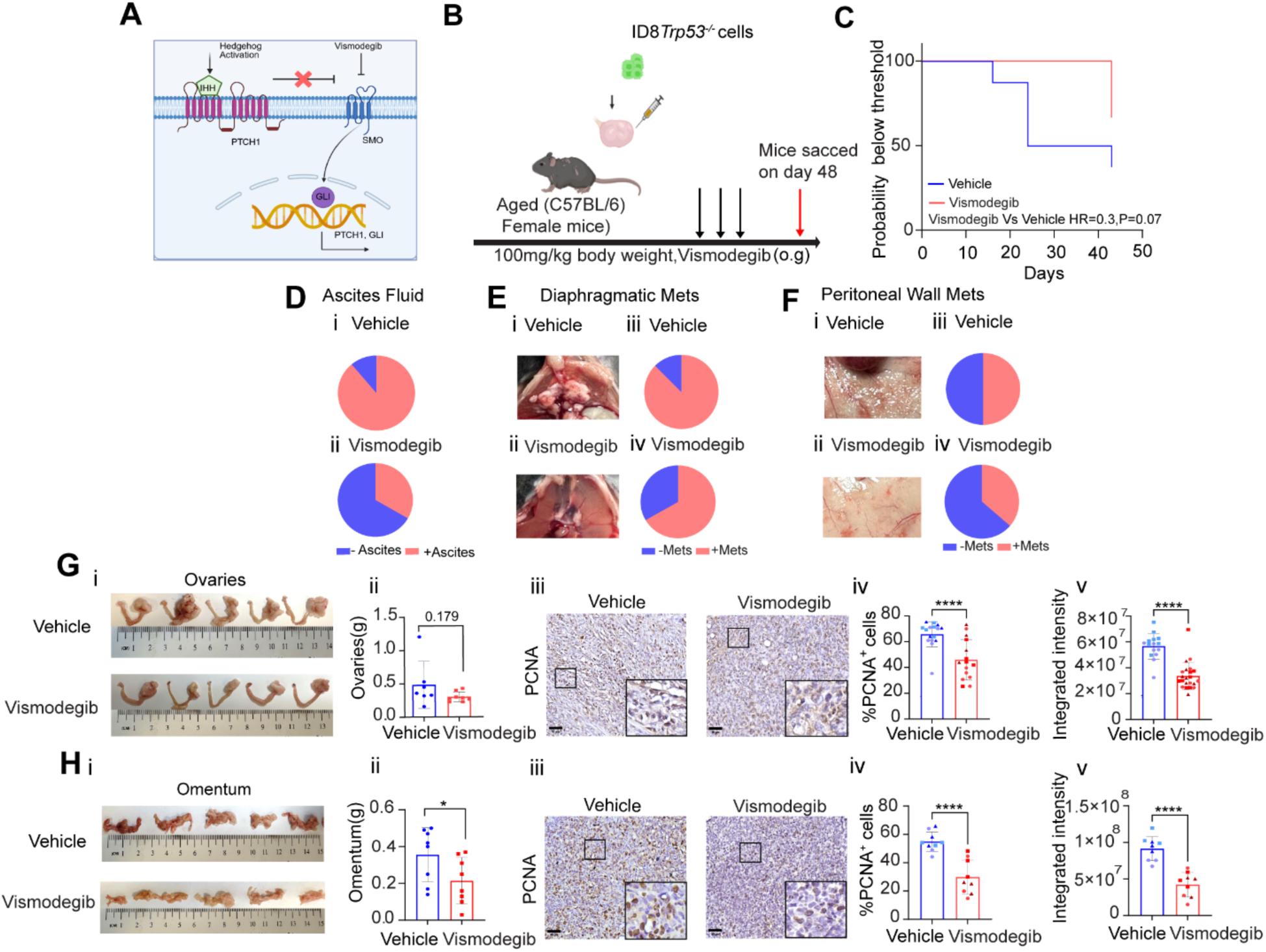
**(A)** Schematic of Hedgehog pathway **(B)** Treatment regimen of Vismodegib post ID8*Trp53^−/−^*cell implantation in the left ovaries of aged mice. **(C)** Probability plot based on abdominal girth changes showing probability of remaining below control derived threshold (n=9). **(D)** Ascites fluid collected from both groups at endpoint, (i) Vehicle (ii) Vismodegib, showing percentage of mice presenting with or without ascites. (**E)** Representative images of diaphragmatic metastatic lesions (mets) (i) Vehicle (ii) Vismodegib, (iii)&(iv) showing percentage of mice with or without diaphragmatic Mets in each group. **(F)** Representative images of mets on peritoneal wall (i) Vehicle (ii) Vismodegib, (iii)&(iv) showing percentage of mice with or without peritoneal wall mets in each group **(G)** (i) Representative images of ovaries and (ii) ovary weights at end points. (iii-v) Representative images of PCNA Immunohistochemistry in ovarian tumor sections with adjacent quantification of %PCNA, positive cells and integrated intensity (n=3). **(H) (i)** Representative images of omentum and (ii) omental weights at endpoint. (iii-v) Representative images of PCNA Immunohistochemistry in omental sections with adjacent quantitation of %PCNA, positive cells and integrated intensity. All data are mean±SEM (n=3), *p<0.05, **p<0.01, ***p<0.001, ***p<0.0001, unpaired t test

### Hedgehog inhibition reduces immunosuppressive macrophages and regulatory T cells in aged tumors

Given the enrichment of Hedgehog signaling in immune cells from aged tumors, we next asked whether the therapeutic effects (Figure 4) were accompanied by changes in immunosuppressive cell populations in ovarian and omental tumors. In ovarian tumors, vismodegib treatment significantly reduced total CD45^+^ cells by 1.85-fold (Supplementary Figure 5A, i) and CD45^+^Arg1^+^ cells by 1.28-fold (Figure 5A), although the latter did not reach statistical significance. Omental tumors showed only marginal changes in these populations (Figure 5B, Supplementary Figure 5A, ii,). While CD8^+^ cells were reduced in both ovarian and omental tumors in aged hosts as compared to young hosts, vismodegib has no effect on this population (Supplementary Figure 5C). Vismodegib has been shown to modulate macrophage polarization in other cancer types ^36,37^. Consistently, CD206^+^ M2 macrophages were 2.4-fold lower in vismodegib-treated ovarian tumors (Figure 5E) but did not reach statistical significance in vismodegib-treated omental tumors (Figure 5F). Flow cytometry analysis of mice ovaries supported these findings, showing trends toward higher M1 (F4/80^+^CD80^+^) and lower M2 (F4/80^+^CD206^+^) macrophages in vismodegib-treated mice (Supplementary Figure 5B, i-v). However, the CD68^+^Arg1^+^ monocytic population was not significantly altered in either ovarian or omental tumors (Figures 5C, 5D). Vismodegib was not cytotoxic to tumor cells in vitro (Supplementary Figure 4C). However, pretreatment of ID8*Trp53^−/−^* tumor cells with vismodegib, reduced CD206 induction in BMDMs cultured with ID8*Trp53^−/−^* conditioned media (Supplementary Figure 5E-H). Vismodegib also lowered CD80 in BMDMs from aged hosts (Supplementary Figure 5E-H) indicating that Hedgehog signaling in tumor cells may contribute to the production of factors that change macrophage activation. HH signaling can also modulate T cell responses in breast cancer^38^. We found that FoxP3^+^ regulatory T cells were markedly reduced in vismodegib-treated mice leading to 2.89-fold reduction in ovarian tumors (Figure 5G, Supplementary Figure 5D) and 1.91-fold lower in omental tumors (Figure 5H, Supplementary Figure 5D) as compared to vehicle treated mice. Together, these findings demonstrate that pharmacologic inhibition of HH signaling in aged hosts suppresses ovarian tumor growth and metastatic spread, accompanied by reductions in CD206^+^ M2 macrophages and FoxP3^+^ regulatory T cells while preserving CD8^+^ T cell frequencies.

**Figure 5.**
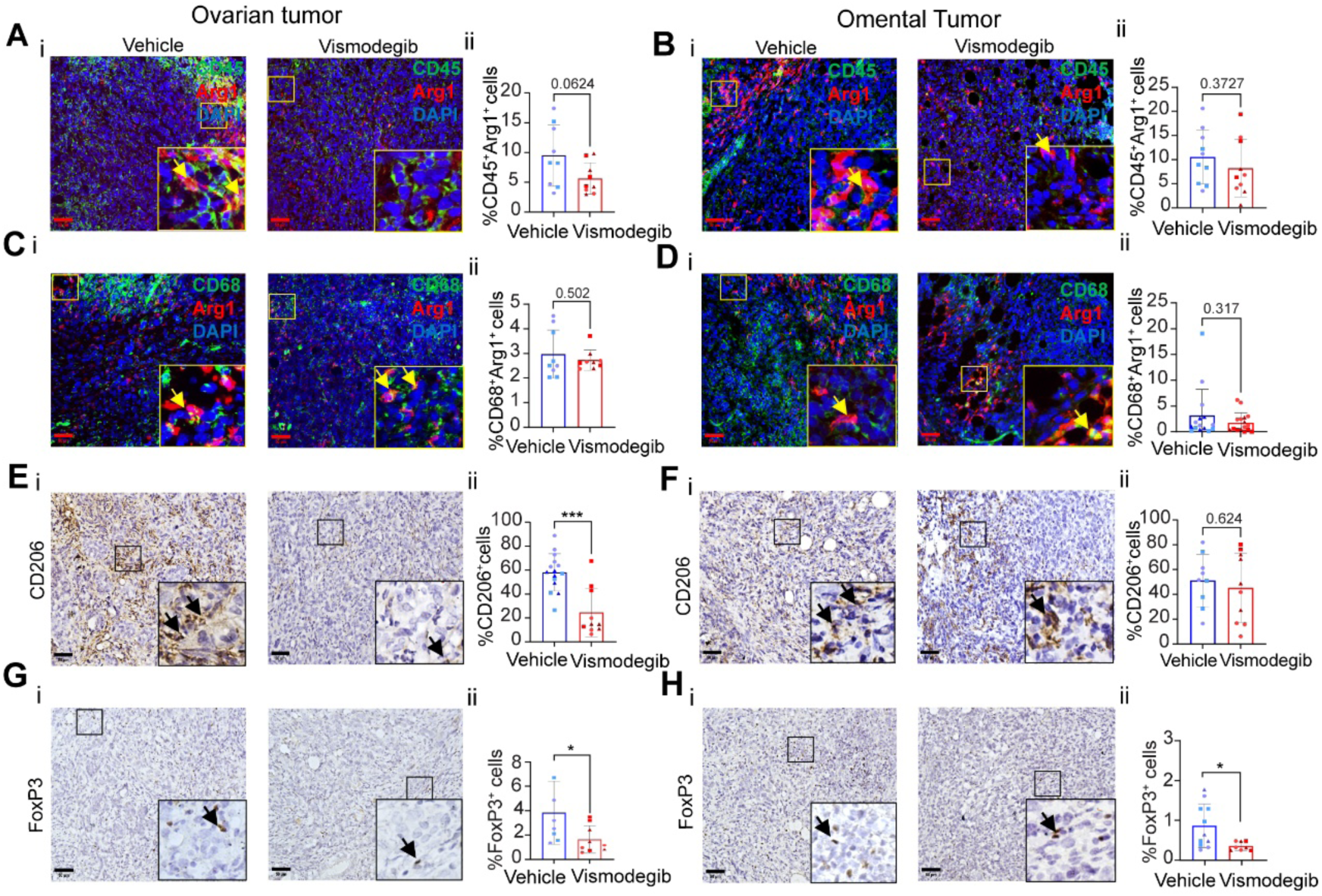
**(A-B)** Representative immunofluorescence images of CD45^+^(green), Arginase-1 (Arg1^+^)(red), and DAPI (blue) in **(A)** ovarian and **(B)** omental tumors from Vehicle or Vismodegib treated mice. Adjacent graphs show percentage of CD45^+^Arg1^+^cells as indicated. **(C-D)** Representative immunofluorescence images of CD68^+^(green), Arg1^+^(red), and DAPI (Blue) in **(C)** ovarian or **(D**) omental tumors from Vehicle or Vismodegib treated mice. Adjacent graphs showing percentage of CD68^+^Arg1^+^cells **(E-F)** Immunohistochemistry of CD206^+^ cells in **(E)** ovarian (**F**) omental tumors. Adjacent graph showing percentage CD206^+^ cells in the omentum. **(G-H)** Representative FoxP3^+^ immunohistochemistry images of (**G)** ovarian or **(H)** omental tumors. Adjacent graph showing percentage Foxp3^+^ cells. Color codes and symbols indicate the number of images analyzed per mouse in each group. All data are presented as mean±SEM (n=3), *p<0.05, **p<0.01, ***p<0.001, ***p<0.0001, unpaired t test.

### Hedgehog pathway activation correlates with immunosuppression in human ovarian cancer

To test the potential relationship between Hedgehog signaling and immunosuppression in human ovarian cancer, we analyzed bulk RNA-sequencing data from TCGA’s ovarian carcinoma samples (TCGA-OV) from cBioPortal. Spearman correlation analysis between Hedgehog pathway genes (*GLI1*, *GLI2*, *PTCH1*, *PTCH2*, *HHIP*, *IHH*, *SHH*) and immune markers revealed selective associations with broadly negative correlations with effector markers (*CD8A*, *CXCL10*, *CCL5*) and more variable associations with immunosuppressive markers (*ARG1*, *MRC1*, *FOXP3*) (Figure 6A). To capture a more composite test of functional balance between immunosuppressive and anti-tumor programs, we correlated ratio variables (*ARG1*/*CXCL10* and *MRC1*/*CD8A*) with a Hedgehog Activation Score (mean z-score of *GLI1*, *GLI2*, *PTCH1*, *PTCH2*, *HHIP*). The *ARG1*/*CXCL10* ratio showed the strongest correlation (ρ=0.450, padj<0.0001), followed by *MRC1*/*CD8A* (ρ=0.283, padj<0.0001) (Figure 6B, Supplementary Figure 6A), indicating that higher Hedgehog pathway activity is associated with a balance favoring suppressive myeloid features over cytotoxic effector programs. A Hedgehog Ligand Score showed similar but weaker correlations with these ratio based immune features (*ARG1*/*CXCL10*: ρ=0.28, padj<0.0001; *MRC1*/*CD8A*: ρ=0.17, padj<0.01) (Supplementary Figure 6B).

**Figure 6.**
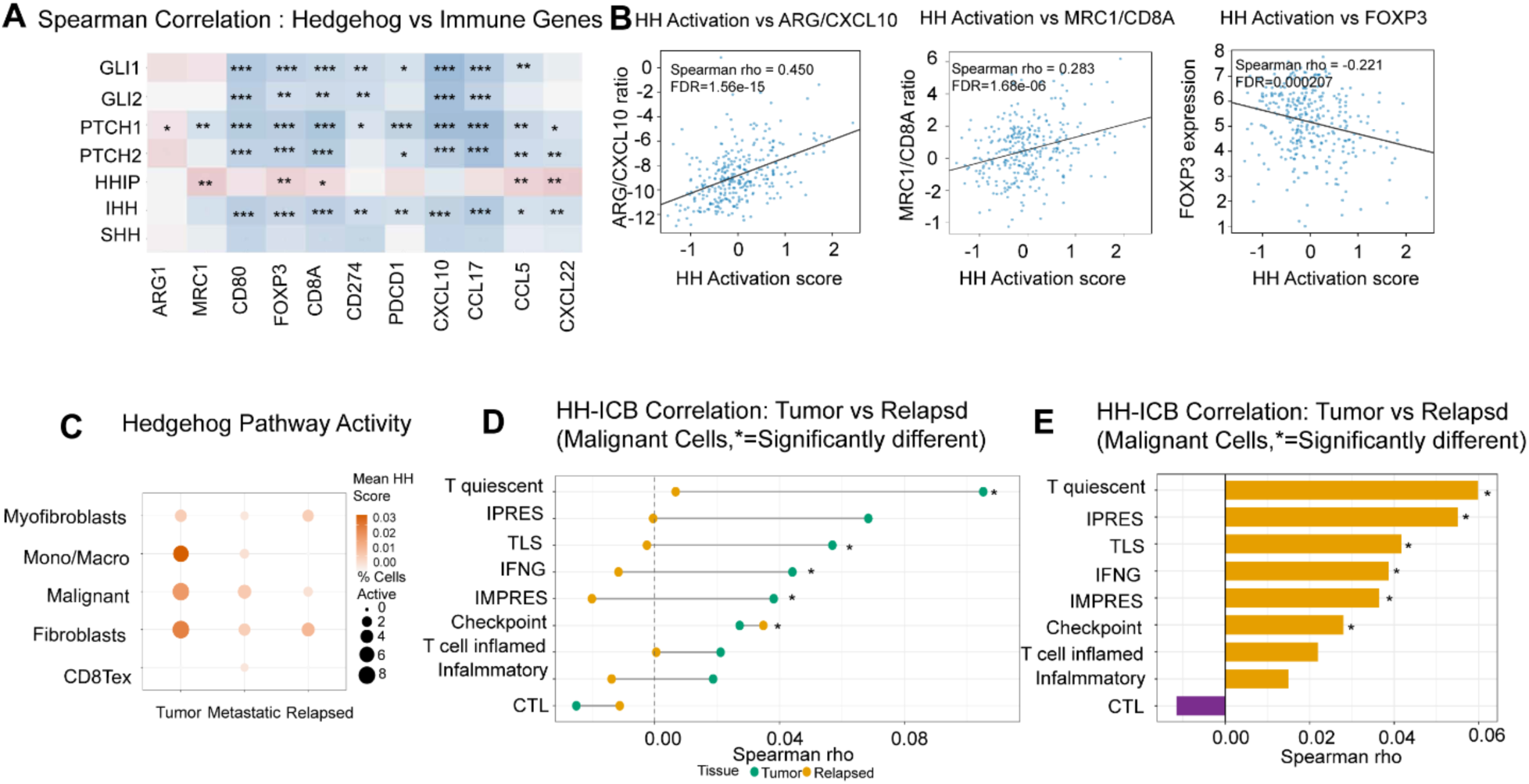
**(A)** Heatmap of Spearman rank correlation coefficients between indicated Hedgehog (HH) pathway genes and selected immune-related genes across TCGA ovarian tumor samples. Color intensity represents direction and magnitude of the correlation, with warm colors indicating positive correlations and cool colors indicating negative correlations. (*p<0.05, **p< 0.01, ***p< 0.001). **(B)** Scatterplots showing selected pairwise relationships between Hedgehog activation score and three selected immune features across TCGA ovarian tumor samples. Each point represents one tumor sample, the solid line indicates the fitted linear trend, and statistical association was assessed using Spearman correlation. **(C)** Dot plot showing Hedgehog pathway activity across five cell types stratified by disease state (primary tumor, metastatic, and relapsed, GSE130000). Dot size represents the percentage of cells with detectable Hedgehog pathway expression (score>0); color intensity represents mean pathway score calculated as the mean log-normalized expression of six detected HH pathway genes (*IHH, PTCH1, PTCH2, HHIP, GLI1, GLI2*) per cell. Data from GSE130000 (n = 11,215 cells). **(D)** Paired dot plot comparing Spearman correlation coefficients between HH pathway score and nine immune checkpoint blockade (ICB) response signatures in primary tumor (teal) versus relapsed (gold) malignant cells. Asterisks denote signatures where the correlation coefficient was significantly different between tumor and relapsed tissue (Fisher’s z-test, Benjamini-Hochberg adjusted *p* < 0.05). **(E)** Bar plot of Spearman correlation coefficients between HH pathway score and nine ICB response signatures across all malignant cells (n = 8,876, GSE130000). (*, Benjamini-Hochberg adjusted p < 0.05).

To extend these findings to single-cell resolution, we analyzed scRNA-seq data from ovarian cancer patients (GSE130000; n=11,215 cells from primary, metastatic, and relapsed tissues across 8 patients). Hedgehog pathway activity, scored as mean expression of six detected genes (*GLI1*, *GLI2*, *PTCH1*, *PTCH2*, *HHIP*, *IHH*), was highest in primary tumors and progressively declined with disease progression (Figure 6C). *PTCH1* and *PTCH2* were most prominently expressed in malignant cells, while *PTCH2* showed highest per-cell expression in monocytes/macrophages. *GLI2* was detected across cell types (Supplementary Figure 6C). Across all malignant cells (n=8,876), HH pathway scores positively correlated with multiple ICB resistance signatures including T-quiescent (ρ=0.060, padj<0.0001), IPRES (ρ=0.055, padj<0.0001), TLS (ρ=0.042, padj<0.001), IFNG (ρ=0.039, padj<0.001), and IMPRES (ρ=0.036, padj=0.001), with no significant association with CTL signatures (Figure 6E). These correlations were significantly stronger in primary tumors compared to relapsed disease based on Fisher’s z- test which confirmed significant attenuation for T-quiescent (Δρ=0.099, padj=0.0001), IPRES (Δρ=0.069, padj=0.007), IMPRES (Δρ=0.059, padj=0.018), Inflammatory (Δρ=0.058, padj=0.018), and T cell-inflamed (Δρ=0.056, padj=0.020) signatures in relapsed relative to primary tumor malignant cells (Figure 6D). Gene-level analysis revealed that *GLI2*, the principal transcriptional effector, specifically correlated with IPRES (ρ=0.037, padj<0.001; Supplementary Figure 6D). No other ICB signature reached significance for *GLI2*, indicating a specific association between active downstream HH transcriptional signaling and features of innate immunotherapy resistance. Gene-level correlation profiles for all HH pathway components across ICB signatures are shown in Supplementary Figure 6E.

Together, these findings across bulk and single-cell human datasets support and extend mouse model observations, demonstrating that Hedgehog pathway activation correlates with an immunosuppressive microenvironment characterized by elevated myeloid suppressor markers, reduced cytotoxic signatures, and ICB resistance programs.

## Discussion

Age is among the strongest risk factors for ovarian cancer, with incidence and mortality rising sharply after menopause, and older patients experiencing poorer outcomes compared to younger patients ^39–41^. While follicle depletion is a hallmark of ovarian aging, whether this change alone accounts for increased tumor susceptibility or whether additional age-associated alterations are required has remained unclear. Using orthotopic implantation into either the ovarian bursa or the peritoneum of two distinct tumor models in young and aged mice, the latter corresponding approximately to human perimenopausal age, we found that aged hosts exhibited significantly higher tumor burden, increased ascites accumulation, and elevated tumor proliferation (Figure 1 A-F). Our data further identified enrichment of Hedgehog signaling in immune cells from aged tumors and demonstrated that pharmacologic inhibition of this pathway reduces tumor burden and immunosuppressive cell populations, suggesting a potential immunomodulatory strategy for post-menopausal patients with ovarian cancer (Figure 2F).

Ovarian aging encompasses far more than follicular decline, including extracellular matrix remodeling, accumulation of senescent stromal and immune cells, and elevated pro-inflammatory cytokines contributing to ‘inflammaging’ ^42^.We used 4-vinylcyclohexene diepoxide (VCD), which selectively destroys primordial and primary follicles, reducing estrogen, progesterone, AMH, and Inhibin A while leaving the young microenvironment otherwise intact ^43^.VCD-treated young mice showed no increase in tumor burden compared to controls, indicating that tumor-promoting effects in aged hosts arise from microenvironmental changes beyond hormonal decline (Supplementary Figure 1D). Prior studies using VCD or ovariectomy in ovarian cancer contexts have yielded variable results. VCD-induced follicle depletion in TgCAG-LS-TAg mice or ovariectomy before SKOV3/OVCAR3 intraperitoneal implantation resulted in slower tumor growth, effects attributed to elevated gonadotropins ^44^. In contrast, VCD treatment followed by carcinogen exposure induced ovarian neoplasms^45^. These studies used intraperitoneal injections or carcinogen-based approaches rather than orthotopic implantation into the ovarian microenvironment, which provides insight into how the aged ovarian niche itself shapes tumor growth. Our orthotopic studies show that follicular depletion is insufficient by itself, but do not identify which specific age-associated changes are necessary or sufficient. The relative contributions of the local ovarian microenvironment versus systemic aging effects, including alterations in circulating immune cells, remain to be determined. Spatial transcriptomics of the tumors revealed that both age groups displayed heterogeneous tumor landscapes with distinct CD45^+^ enriched and depleted regions; the pathway signatures defining these regions differed substantially between age groups. In young host tumors, CD45^+^ depleted regions showed hallmarks of metabolic reprogramming while CD45^+^ enriched regions exhibited epithelial-mesenchymal transition (EMT) signatures. Emerging evidence indicates that EMT can serve as a tumor escape mechanism in response to anti-tumor immune pressure, enabling tumor cells to evade T cell-mediated killing through MHC class I downregulation and immune checkpoint upregulation (Figure 2C, i) ^46–48^. The presence of EMT signatures in immune-enriched regions of young tumors, alongside anti-tumor chemokines such as *CXCL10*, suggests that tumors in young hosts may face greater immune pressure and utilize EMT as an adaptive response. In contrast, aged host tumors showed MYC targets and cholesterol homeostasis enrichment in CD45^+^ depleted regions, and inflammatory response signatures in CD45^+^ enriched regions (Figure 2C, ii). Notably, while tumors from both young and aged hosts exhibited inflammatory hallmarks in immune-enriched regions, the genes driving these signatures were entirely non-overlapping (Figure 2D). This distinction has functional implications such as with *Cxcl10*, enriched in young tumors, which is associated with anti-tumor immunity and a favorable prognosis in ovarian cancer ^49^, whereas chemokines enriched in aged tumors, including *Ccl5, Ccl22,* and *Ccl17* promote immunosuppression through regulatory T cell recruitment and are associated with chemoresistance and poor response to immunotherapy ^50–53^. These findings reveal that similarly labeled inflammatory signatures obscure fundamentally different immune programs indicating that tumors from young hosts appear to face active anti-tumor immunity necessitating escape mechanisms, whereas tumors from aged hosts exhibit signatures consistent with an already established immunosuppressive microenvironment. B cell-associated genes (*Gsk3a*, *Txndc5*, *Itga5*) and Jchain were similarly enriched in tumors from young hosts (Figure 2G, Supplementary Figure 2B, i-ii), with B cells contributing to anti-tumor immunity in HGSOC through antibody-mediated tumor recognition and ADCC, antigen presentation, tertiary lymphoid structure formation, and polymeric IgA transcytosis ^54^. Whether EMT-targeted approaches, restoration of humoral immunity, or other strategies could enhance anti-tumor responses in younger patients, or whether reversing the immunosuppressive chemokine milieu could benefit older patients, warrants further investigation.

A second seemingly paradoxical finding from the spatial transcriptomic analysis was the KEGG enrichment of natural killer cell cytotoxicity pathways in CD45^+^ enriched regions of aged tumors (Supplementary Figure 2A, iv), which initially appears at odds with the broader immunosuppressive phenotype. However, NK cells in the tumor microenvironment can exhibit intact transcriptional cytotoxicity signatures without effective tumor killing - a documented pattern of NK functional exhaustion in ovarian cancer. Factors enriched in the aged TME, including Tgf-β, Pge2, and adenosine, impair NK degranulation independent of gene expression, and chronic IFN-α/IFN-γ exposure (Figure 2D, F) can transcriptionally drive cytotoxicity genes via STAT signaling while paradoxically inducing NK exhaustion through inhibitory checkpoint upregulation, a hallmark of inflammaging. Consistent with this dissociation, CIBERSORTX deconvolution (Figure 2E) revealed comparable or slightly reduced NK cell percentages in aged tumors, indicating that transcriptional pathway activity does not equate to increased effector cell abundance or function. Similarly, the IFN-α and IFN-γ signatures observed in aged tumor CD45^+^ cells (Figure 2F) require contextual interpretation. While acute IFN responses promote anti-tumor immunity through dendritic cell activation, MHC-I upregulation, and CD8^+^ T cell priming, chronic IFN exposure drives immunosuppression via checkpoint molecule upregulation (*PD-L1*, *IDO*, *LAG3*), M2 macrophage polarization through STAT signaling, and T cell exhaustion. The IFN signatures we observe coincide with elevated CD206^+^ M2 macrophages, FoxP3^+^ Tregs, reduced CD8^+^ T cells, and accelerated tumor growth, a context consistent with chronic rather than acute IFN signaling, characteristic of the inflammaging state associated with physiological aging.

Consistent with our transcriptomic findings, aged host tumors exhibited significantly higher infiltration of immunosuppressive populations across both tumor models, including CD45^+^Arg1^+^ cells, which were significantly higher in aged ovarian and omental tumors compared to young hosts (Figures 3). This enrichment was driven by CD68^+^Arg1^+^ monocytes and CD206^+^ tumor-associated macrophages (TAMs), both of which were significantly elevated in aged tumors. CD206^+^ TAMs are established drivers of immunosuppression in ovarian cancer and are associated with poor prognosis, therapy resistance, and tumor progression^24,55,56^. The balance between CD80^+^ and CD206^+^ macrophage phenotypes is critical in shaping the local immune milieu, and our finding of a higher F4/80^+^CD206^+^/F4/80^+^CD80^+^ ratio in ascites from aged mice further supports a shift toward immunosuppressive macrophage polarization in aged hosts (Supplementary Figure 3B, iv, v).

More broadly, recent single-cell studies of healthy aged ovaries have shown expanded double-negative T cells, increased tissue macrophages, and reduced leukocyte chemoattractant ^14,57^.The immunosuppressive phenotype we observe in aged tumor-bearing ovaries likely reflects both these baseline aged immune changes and tumor-driven amplification, though our design does not formally separate these contributions. The macrophage landscape in HGSOC is however heterogeneous, including tissue-resident peritoneal macrophages, monocyte-derived populations, lipid-associated macrophages (LAMs), and spatially distinct tumor-core and invasive-edge subsets. Our CD206 and Arginase 1 (Arg1) phenotyping captures macrophages with pro-tumor immunosuppressive features but does not resolve these finer subsets; single-cell or spatial transcriptomic approaches will be needed to dissect subpopulation-specific responses to Hedgehog inhibition.

In parallel, FoxP3^+^ regulatory T cells (Tregs) were more abundant in tumors from aged hosts across both ovarian and omental sites (Figures 3). Tregs inhibit effective anti-tumor immunity and have been associated with poor survival, specifically in ovarian cancer, compared to other malignancies^58^. Aged hosts had higher double-positive MDSCs: CD11b^+^Ly6G^+^Lyc6C^+^, which are particularly immature and highly activated, suppressive populations ^59^ (Supplementary Figures, 3B, xv). The concurrent enrichment of M2 macrophages and regulatory T cells in tumors from aged hosts suggests that these populations may cooperate to establish an immunosuppressive microenvironment that permits accelerated tumor growth. Whether this immunosuppressive state reflects intrinsic properties of aged immune cells, tumor or stroma mediated reprogramming of infiltrating immune cells, or altered recruitment patterns driven by the aged host remains to be determined. A related question is whether the observed phenotype simply reflects baseline differences in immune populations between the ovary and the peritoneal cavity, independent of host age. Human peritoneal fluid is enriched for macrophages, T cells, dendritic cells, NK cells, and B cells^60,61^; the mouse peritoneal cavity is dominated by embryonically-derived large peritoneal macrophages that are replaced by monocyte-derived small peritoneal macrophages during inflammation^62^ ^63^; and ovarian macrophages function primarily in tissue remodeling and T cell surveillance^64^. Despite these compositional differences, the same age-associated immunosuppressive profile of elevated CD206^+^ TAMs, FoxP3^+^ and reduced CD8^+^ T cells, was observed in both ovarian and omental tumors across regardless of delivery route (i.b versus i.p) (Supplementary Figures 3A, 3C), arguing that physiological aging creates a systemically immunosuppressive environment that overrides baseline compartmental differences. This raised the question of which molecular pathways might be implicated in this age-associated immune shift.

The Hedgehog pathway, while best known for its roles in embryonic development and tissue homeostasis, has emerged as an important modulator of immune function in the tumor microenvironment ^65,66^. Recent studies in breast cancer have begun to elucidate how Hedgehog signaling shapes immunosuppressive cell populations. Hedgehog signaling regulates macrophage metabolism and polarization, promoting the M2 phenotype through altered O-GlcNAcylation and mitochondrial dynamics ^26^. Additionally, Hedgehog signaling promotes Treg differentiation and activity, and its inhibition can drive Treg-to-Th17 conversion through metabolic rewiring ^38^. Consistent with this microenvironment-centric mechanism, vismodegib was minimally cytotoxic to murine tumor cells viability (Supplementary Figure 4) arguing that the reduced tumor proliferation observed in vivo (Figure 4) reflects effects on the host tumor microenvironment rather than direct cytotoxicity. The mechanistic insights from breast cancer models suggest potential pathways through which Hedgehog activation in ovarian tumors from aged hosts could sustain immunosuppressive M2 macrophages and Tregs. Our data demonstrates correlation, and whether Hedgehog activation is directly driving immunosuppressive cell polarization in aged ovarian tumors, or whether it represents a consequence of the altered aged microenvironment, remains to be determined.

Our flow panels do not directly address the role of Th17 cells that can increase with age in postmenopausal women and contribute to tumor progression through recruitment of immunosuppressive populations including MDSCs and enhancement of angiogenesis^67^. We note that IL-6/STAT3 signaling, a known Th17 driver, was enriched in CD45^+^ regions of young host tumors in our spatial data (Figure 2C), consistent with active immune engagement in young hosts. Separately, Hedgehog signaling has been shown to regulate Treg-to-Th17 conversion in breast cancer, with vismodegib shifting cells away from a Treg phenotype ^38^.The FoxP3^+^ Treg reduction we observe with vismodegib may therefore reflect both direct effects on Treg differentiation and partial Treg-to-Th17 conversion, with formal Th17 characterization in aged hosts representing an important future direction.

Beyond Th17 considerations, our broader vismodegib results showed delayed disease progression with decreased infiltration of CD206^+^ M2 macrophages and FoxP3^+^ Tregs, while CD68^+^Arg1^+^ monocytes were not significantly affected and CD8^+^ T cell frequencies were preserved (Supplementary Figure 5). This pattern of preserved CD8^+^ T cell infiltration in aged ovarian tumors contrasts with a prior report ^68^, where Smo inhibition diminished CD8^+^ migration in colorectal cancer and BCC through a noncanonical Smo–RhoA mechanism. The Hedgehog pathway activation we observe specifically within immune populations of aged tumors (Figure 2) may favor effects on pathway-active immunosuppressive cells rather than uniform impairment of CD8^+^ migration, suggesting that HH pathway inhibition effects are tumor-type and microenvironment-context dependent.

Our preclinical observation of vismodegib efficacy in aged hosts contrasts with a phase II clinical trial in which vismodegib failed to improve progression-free survival as maintenance therapy in ovarian cancer patients in second or third complete remission ^69^. Several factors may account for this discrepancy. First, patients in that trial were not selected based on age or profiled for Hedgehog pathway activation, and Hedgehog ligand expression was detected only in 13.5% of archival tissues^69^. Our data suggest Hedgehog signaling is specifically enriched in the aged tumor microenvironment, and patients without pathway activation may derive little benefit. Our analysis of human OCs (Figure 6) demonstrates that hedgehog pathway activation correlates with specific immunosuppressive signatures providing a framework for biomarker based patient selection in future trials. Second, the clinical trial employed vismodegib as maintenance therapy after achieving remission, whereas our intervention began early after tumor implantation when the immunosuppressive microenvironment was actively developing. Our single-cell re-analysis of human ovarian cancer further supports this point, showing that Hedgehog pathway activity is highest in primary tumors and progressively attenuated through metastatic and relapsed disease (Figure 6C, 6D), suggesting that vismodegib maintenance therapy administered to patients in remission may have been initiated at a disease stage when the pathway is no longer a significant driver of immunosuppression. Perhaps most importantly, the immunomodulatory effects we observe suggest that Hedgehog inhibition may be best suited not as monotherapy but in combination with immune checkpoint inhibitors. By reducing M2 macrophages and Tregs, Hedgehog inhibition could potentially render tumors more responsive to checkpoint blockade, a combinatorial strategy currently being evaluated in a phase II trial of vismodegib plus atezolizumab in ovarian cancer (NCT05538091), and warranting further investigation in age and/or pathway-stratified cohorts, particularly in post-menopausal patients whose tumors may be most dependent on this pathway.

Some limitations of our study warrant acknowledgment. The TCGA-OV cohort skews toward older, advanced-stage patients with heterogeneous mutational backgrounds, including BRCA1/2 alterations that are more frequent in younger HGSOC and influence the immune microenvironment. The scRNA-seq analysis validation is drawn from patients without indicated BRCA alterations, but a definitive age-stratified human analysis will require BRCA-stratified cohorts. Additionally, while host mice were aged, the implanted tumor cells originated from young mice and therefore do not model age-associated alterations in tumor cell-intrinsic biology. Future studies using aged BRCA1/2-mutant cell lines in aged hosts, or cell lines/genetic models derived from aged hosts will be needed to distinguish host versus tumor-intrinsic age effects and the age-by-genotype interaction. Comprehensive phenotyping of stromal, vascular, and additional immune populations (e.g., T cell functional characterization) including scRNA-seq of young and aged tumors with and without Hedgehog inhibition will also be needed to fully characterize the effects of Hedgehog inhibition on the tumor microenvironment. Definitive proof that CD206^+^ macrophages and FoxP3^+^ Tregs are necessary for age-associated tumor progression will require in vivo depletion experiments in aged tumor-bearing mice which represents an important future direction.

Together, our data indicates that physiological aging creates an immunosuppressive ovarian tumor microenvironment characterized by CD206^+^ tumor-associated macrophage and FoxP3^+^ regulatory T cell infiltration, with Hedgehog signaling emerging as a potential mediator. The distinct immune landscapes in young versus aged tumors, active immune engagement and potential for EMT-mediated escape in young hosts versus pre-established immunosuppression in aged hosts suggest that age-stratified therapeutic approaches may be warranted. Most immediately, our data provides rationale for clinical investigation of Hedgehog inhibitors in combination with immune checkpoint blockade, with patient selection based on age and tumor Hedgehog pathway activation status.

## Acknowledgements

Funding for this work was provided in part by NIHR01CA219495 and O’Neal Invests grant to Mythreye Karthikeyan (KM). This study was supported by National Institutes of Health (NIH) award NIHR00AG068309 an AFAR Grant for Junior Faculty awardee to Daniel Tyrrell (DJT). The funders had no role in study design, data collection and analysis, decision to publish, or preparation of the manuscript. We want to acknowledge the UAB flow cytometry core facility, (AI027767), O’Neal Cancer Center, P30CA013148, and shared instrument grant S100D032296. We also acknowledge the Preclinical Imaging Shared Facility (P30CA013148, 1S10OD021697), High Resolution Imaging Facility and the Pathology Core Research Lab at UAB for assistance with processing the histological specimens. Schematics were made using Biorender (licensed agreement to KM and UAB). We thank Amr Ismail and Emily O’Brian for technical assistance, Dr. Arend for sharing cell lines and Dr. Darshan Shimoga Chandrashekar for helpful discussions.

## Author contributions

Conceptualization: KM, AK. Investigation: AK, CM. MHE, RR, KS, MM, LMQ, DJT, LAS, CMM, RR, AK, LQM, MVN, SS, FM, MC and SA. Analysis: KM, CM, AK, KS, SA Reagents: AK, KF, DJT, LAS Resources/Supervision: KM, CM, DT, LAS. Writing: original draft: AK, KM Writing: review & editing: all authors. Funding acquisition as described in acknowledgements and project administration: KM.

## Data availability statement

GeoMx data are available at doi: 10.17632/j4f8r6z3b3.1. Additional data are available from the corresponding author upon reasonable request. Code for the reanalysis of single-cell GSE130000 is deposited at GitHub (https://github.com/mythreyelab/Hedgehog-ovarian-scRNAseq) and archived at Zenodo (DOI: https://doi.org/10.5281/zenodo.20542282). Code for the TCGA-OV bulk-expression analysis is deposited at GitHub (https://github.com/mythreyelab/Hedgehog-ovarianimmune-TCGA)and archived at Zenodo (DOI: https://doi.org/10.5281/zenodo.20542584). DOIs are listed in the Key Resources Table.

## STAR METHOD

Detailed methods are provided in the online version of this paper and include the following:

- KEY RESOURCES TABLE
- EXPERIMENTAL MODEL AND STUDY PARTICIPANT DETAILS

- Cell lines.
- Animals.
- METHOD DETAILS

- Establishment of follicle-depleted ovaries.
- Intrabursal implantation of ovarian cancer cells
- Intraperitoneal implantation of ovarian cancer cells.
- In vivo vismodegib treatment.
- In vitro vismodegib treatment.
- Endpoint tumor burden measurements.
- Time-to-Threshold Analysis (Fixed).
- Time-to-Threshold Analysis (Control-Derived).
- Spatial Nanostring GeoMx analysis.
- Hedgehog pathway analysis in human data.
- Immunohistochemistry.
- Immunofluorescence.
- Flow cytometry.
- qRT-PCR for Hedgehog pathway.
- Bone marrow-derived macrophage (BMDMs) polarization.
- QUANTIFICATION AND STATISTICAL ANALYSIS

### EXPERIMENTAL MODEL AND STUDY PARTICIPANT DETAILS

#### Cell lines

PPNM (*p53^−/−l^ ^R172H^ Pten^−/−^Nf1*^−/−^*Myc^OE^*) cells were kindly provided by Dr. Robert A Weinberg, Whitehead Institute for Biomedical Research, Cambridge, MA, USA, through an MTA and were cultured as described here ^21^. For ID8*Trp*^53^*^−/−^* Luc, ID8*Trp53^−/−^*cells ^70^ were transduced with Luc FUW-firefly luciferase-EGFP (FUW-FFLuc-eGFP) lentiviral construct gene, generously gifted by Dr. Stewart, Department of Cell Biology and Physiology, Washington University School of Medicine, St Louis, to generate ID8*Trp53^−/−^* Luc cells as described^71^. Both cells lines were authenticated by providers. ID8*Trp53^−/−^* cells were cultured in Dulbecco’s modified Eagle’s medium supplemented with 4% fetal bovine albumin, 100U penicillin, and streptomycin, 5 μg/mL of insulin, 5 μg/mL of transferrin, and 5 ng/mL of sodium selenite (41400045, Gibco). Both cell lines were maintained at 37 °C with 5% CO_2_ in a humidified incubator, routinely checked for mycoplasma, and experiments were conducted within 3–6 passages, depending on the cell line.

#### Animals

All animal procedures were conducted in accordance with Institutional Animal Care and Use Committee (IACUC) guidelines at the University of Alabama at Birmingham (UAB), USA. C57BL/6J (000664; The Jackson Laboratory, Farmington, CT, USA,) or C57BL/6N (National Institute of Aging, Bethesda, MD, USA)) nulliparous female mice aged 6 or 65 weeks, were used in this study. All mice were housed in the Wallace Tumor Institute animal facility at UAB with food and water ad libitum in groups of 5 mice per cage at 24°C under a 12-hour light/dark cycle. Only female mice were used in this study as ovarian cancer is a sex specific malignancy. Consistently, sex based comparisons were not applicable.

### METHOD DETAILS

#### Establishment of follicle-depleted ovaries

Young female mice (6 weeks old) were randomly divided into two groups (n=5 per group): 4-vinylcyclohexene diepoxide (VCD, 94956, Sigma-Aldrich)) and Sesame oil (S3547, Sigma Aldrich) treated. The VCD group received intraperitoneal injections at 160 mg/kg in Sesame oil for 15 consecutive days^72^. Control mice received sesame oil alone; following VCD treatment, mice were aged until postnatal day 60, at which point ovaries were collected for histological analysis for follicle counts. Anti-müllerian hormone (AMH) and inhibin A (Supplementary Table C) levels were quantified in ovaries post cDNA synthesis using reverse transcriptase kit (Cat. no. 95047-500, Quantabio) qRt-PCR using iTaq SYBR Green Super mix (Cat no. 1725125, Bio-Rad), Rpl was used as house keeping gene. Relative expression was calculated using the 2−ΔΔCt method, normalized to vehicle-treated controls.

#### Intrabursal implantation of ovarian cancer cells

2x10^6^ ID8*Trp53*^−/−^ or PPNM cells suspended in 8µl phosphate-buffered saline were injected into the bursa of the left mouse ovary. Mice were anesthetized prior to implantation using 2.5% isoflurane mixed with 2L/min oxygen and administered buprenorphine 0.07 mg/kg as analgesic pre- and post-surgery for up to 48 hours following institutional guidelines.

#### Intraperitoneal implantation of ovarian cancer cells

2x10^6^ PPNM cells suspended in 100µl phosphate-buffered saline were injected into the intraperitoneal cavity of female young (6-8weeks old) and aged (60-65 weeks old) mice.

#### In vivo vismodegib treatment

Aged female mice (65 weeks old) were gavaged with 100µl (100mg/Kg) of Vismodegib (GDC-0449) (Cat no. S1082, Selleckchem) or DMSO as vehicle, three times a week as described previously^26^ after 48 hours of ID8*Trp53*^−/−^ cell injection in the ovaries. This dose and schedule were selected based on prior C57BL/6 mouse studies demonstrating efficacy without toxicity^38,73,74^. Body weight was monitored throughout the treatment. Mice were also assessed weekly for activity, grooming, and clinical signs including hunched posture, piloerection, and diarrhea; none were observed in either group. Mice were closely monitored for post-surgical recovery over 5 days, and tumor growth was tracked every 10 days using whole-body bioluminescence imaging (BLI) on a Perkin Xenogen IVIS Imaging System following intraperitoneal injection of luciferin (50 mg/kg).

#### In vitro vismodegib treatment

ID8*Trp53^−/−^*and PPNM cells were seeded at 1,000 cells/well in 96-well plates and allowed to adhere overnight. Cells were treated with vismodegib at a range of concentrations ranging from 50 μM/ml to 0.2 μM/ml or DMSO vehicle (0.25 % v/v) for 48 hours. Cell viability was assessed using Sulforhodamine B (SRB) assay as described previously^75^. Briefly, cells were fixed in 10% Trichloroacetic acid (200 µl per well) for 10 min at 4°C. followed by washing with distilled water three times for 2 min each. Sulforhodamine B in 1% glacial acetic acid was added at 100 µl/well and incubated at room temperature for 10 min, followed by washing with 1% glacial acetic acid three times, 1 min each. Plates were left to air-dry overnight, 10 mM Tris buffer (100 µl per well) was added, and absorbance was read at 570 nm using a Synergy H1 microplate reader (Biotek, Winooski, VT, USA). IC50 was calculated using nonlinear regression using GraphPad-Prism 9.0. Experiments were performed in technical triplicate and biological triplicate.

#### Endpoint tumor burden measurements

At the experimental endpoint, ovaries, omentum, and peritoneal/mesenteric tumor deposits were collected and weighed individually. Ascites fluid was aspirated from the peritoneal cavity and quantified by volume. Total tumor burden was reported as the sum of ovarian, omental, and peritoneal/mesenteric weights

#### Time-to-Threshold Analysis (Fixed)

Disease progression in comparing young versus old mice was quantified using a time-to-threshold framework in which the event was defined as the first attainment of a pre-specified biological disease burden. For peritoneal dissemination studies, the threshold was defined a priori as ≥2.77 cm² peritoneal spread or a corresponding abdominal girth criterion reflecting maximum tolerable tumor burden. The 2.77 cm² peritoneal spread threshold was empirically defined as the disease burden at which mice consistently met institutional humane endpoint criteria (reduced mobility, hunched posture, and other IACUC-defined morbidity indicators), providing a proxy for disease severity. For each mouse, time to reach this threshold was recorded as the event. Kaplan–Meier curves were generated and compared using the log-rank test, with hazard ratios estimated by Cox proportional hazards modeling.

#### Time-to-Threshold Analysis (Control-Derived)

In studies using abdominal girth as the progression metric, a control-derived reference threshold was employed. The threshold was defined exclusively from the control cohort as the mean change in abdominal girth from baseline to endpoint. This control-derived benchmark was then applied uniformly to both control and treatment groups. For each mouse, the time to reach this benchmark was scored as the event. Kaplan Meier curves were generated and compared using the log-rank test, with hazard ratios estimated using Cox proportional hazards regression. Abdominal girth was measured weekly at the widest point of the abdomen using a flexible measuring tape by an investigator blinded to treatment assignment.

#### Spatial Nanostring GeoMx analysis

RNA profiling of ovarian tumors from young and aged mice was performed using the mouse Whole Transcriptome Atlas probes using GeoMx™ DSP Nanostring technology per manufacturer’s guidelines. In brief, paraffin-embedded 5µm ovarian sections were baked at 60°C for 1 hour. They were stained with labeled Goat anti-mouse CD45 (AF114, 1:100, Novus Biologicals) primary antibody labeled with FITC using FITC Conjugation Kit (ab188285, Abcam) and Rat anti-mouse CK8 primary antibody (Sc8020, 1:100, Santa Cruz Biotechnology) labeled with APC using APC Conjugation Kit (ab201807, Abcam) for immune and tumor cells detection, respectively. Following incubation with primary antibodies for 2 hours at room temperature, nuclei were stained with SytoTM82, an Orange fluorescent Nucleic acid stain (S11363, 1:1000, Sigma-Aldrich). Regions of interest (ROIs) were selected using a combination of immunofluorescent staining and H&E staining. ROIs were classified as general areas to obtain an unbiased overview of the microenvironment such as tumor cells (CK8^+^) or immune cells (CD45^+^). CD45^+^ enriched regions were defined as ROIs containing >50 CD45^+^ cells, while CD45^+^ depleted regions contained <10–15 CD45^+^ cells, based on automated cell counting from the immunofluorescence channel. These thresholds were selected to capture distinct immunological environments while ensuring sufficient tissue area for GeoMx profiling. Upon ROI collection on the GeoMx Digital Spatial Profiler (DSP), libraries were prepared and sequenced on the Illumina NovaSeq instrument (UAB, Genomics core facility). FASTQ files were uploaded to the Base Space Illumina hub and converted to digital count conversion (dcc) files using the GeoMx® NGS Pipeline (v2.0.21) on Illumina DRAGEN. The analysis was performed in R following the manufacturer’s analysis code (https://www.bioconductor.org/packages/release/workflows/vignettes/GeoMxWorkflows/inst/do <u>c/GeomxTools_RNA-NGS_Analysis.html</u>). ROIs with less than 80% sequencing alignment or less than 50% sequencing saturation were removed from further analysis. The Grubbs outlier test was used to identify outlier probes. The limit of quantification (LOQ) was defined as two standard deviations above the geometric mean of the negative probes. ROIs were then divided into general areas, and CD45^+^ immune cell genes below LOQ in at least 10% of ROIs were removed from further analysis. Normalization was performed using a signal-based quartile normalization method, where individual counts are normalized against the 75th percentile of signal from their ROI. Group comparisons were done using a mixed linear model as previously described. Normalized counts can be accessed here: doi: 10.17632/j4f8r6z3b3.1. Normalized gene counts were used for Gene Set Enrichment Analysis (GSEA) using GSEA 4.3.3 (Broad Institute, UCSD, San Diego)^76^, and immune cell abundances analysis CIBERSORTX was used after converting gene IDs from mouse to human on SynGO (Synaptic Gene Ontologies and annotations) ^23,77^. Immune infiltration association with ovarian cancer patient survival was performed using Timer 2.0^78^. Heat maps were generated using Heatmapper2 ^79^ and KEGG pathway analysis was performed using the SRPlot platform ^80^.

#### Hedgehog pathway analysis in human data

##### Datasets

For bulk transcriptomic analysis, mRNA expression data from The Cancer Genome Atlas ovarian cancer cohort (TCGA-OV) were obtained from cBioPortal (*data_mrna_seq_v2_rsem.txt*; n = 300 tumor samples)^81–83^. Expression values are RSEM-normalized. Rows with missing Hugo gene symbols or duplicate gene symbols (retaining the first occurrence) were excluded prior to analysis. For single-cell transcriptomic analysis, we used the publicly available scRNA-seq dataset GSE130000 (n = 11,215 cells from eight patient samples spanning three disease states: primary tumor [patients P1–P4; n = 5,215 cells], metastatic [M1–M2; n = 1,474 cells], and relapsed [R1–R2; n = 4,526 cells]). Cells were annotated into five major lineages - malignant cells (n = 8,876), CD8+ exhausted T cells (CD8Tex; n = 372), fibroblasts (n = 884), monocytes/macrophages (n = 259), and myofibroblasts (n = 824) - with annotations obtained from the TISCH2 database metadata. The expression matrix was stored in HDF5 format and loaded using the hdf5r and Matrix R packages. Gene expression values were normalized to counts per 10,000 (CP10K) and log-transformed using the log1p function, consistent with standard scRNA-seq normalization approaches^84^.

##### Hedgehog pathway gene panel

Hedgehog pathway activity was assessed using a panel of seven canonical genes comprising two ligands (*SHH, IHH*), two receptors (*PTCH1, PTCH2*), one hedgehog-interacting protein (*HHIP*), and two GLI transcription factors (*GLI1, GLI2*). This panel was applied in the TCGA-OV bulk dataset across all 300 tumor samples. In the GSE130000 scRNA-seq analysis, *DHH* was additionally queried but was not detected above background, nor was *SHH*; the six detected genes (*IHH, PTCH1, PTCH2, HHIP, GLI1, GLI2*) were used for scoring.

##### Hedgehog pathway scoring

In the TCGA-OV dataset, pathway-level scores were computed at the sample level. The Hedgehog Activation Score was defined as the mean z-score of log2-transformed expression values for *GLI1, GLI2, PTCH1, PTCH2*, and *HHIP*. A Hedgehog Ligand Score was defined separately as the mean z-score of log2-transformed *IHH* and *SHH*. Gene-level correlations were additionally examined directly from the processed expression matrix. In GSE130000, a per-cell HH pathway score was calculated as the mean log-normalized (CP10K + log1p) expression across the six detected HH genes. Primary HH–immune correlation analyses were conducted in malignant cells (n = 8,876); gene-level analyses (individual HH gene expression versus ICB signatures) were performed across all cells (n = 11,215).

##### Immune features

In the TCGA-OV analysis, pairwise Spearman correlations were computed between HH pathway genes or scores and a panel of 11 immune-associated genes: *ARG1, MRC1, CD80, FOXP3, CD8A, CD274, PDCD1, CXCL10, CCL5, CCL17*, and *CCL22*. Immune and chemokine markers were selected based on established roles as immunosuppressive or immune-regulatory mediators in human high-grade serous ovarian cancer, together with features highlighted by spatial transcriptomic inflammatory-hallmark analyses in the murine model. To capture the functional balance between immunosuppressive and cytotoxic effector programs, two composite ratio metrics were computed: ARG1/CXCL10 and *MRC1/CD8A.* Ratios were calculated as log2((numerator + 1) / (denominator + 1)) to stabilize values at low expression levels.

In the GSE130000 scRNA-seq analysis, HH pathway scores were correlated with nine published immune checkpoint blockade (ICB) response signatures: Checkpoint (inhibitory immune checkpoint genes), CTL (cytotoxic T lymphocyte effector genes; ^85^, IFNG (interferon-gamma 6-gene signature)^86^, IMPRES (Immune Predictive Score)^87^, Inflammatory (inflammatory cytokine and chemokine genes)^88^, IPRES (Innate anti-PD1 Resistance) ^89^, T cell-inflamed (18-gene T cell-inflamed gene expression profile) ^86^, TLS (tertiary lymphoid structure signature)^90^, and T-quiescent (quiescent T cell markers)^91^. Each signature was scored per cell using the mean log-normalized expression approach described above.

##### Software

All analyses were performed in R (version 4.5.3). Spearman rank correlations were computed with ties handled via an asymptotic approximation; multiple-testing correction used the Benjamini–Hochberg procedure. The TCGA-OV figures were generated using base R graphics and exported as PNG and PDF files. The scRNA-seq analysis additionally used: hdf5r for HDF5 file access, Matrix for sparse matrix operations, dplyr and tidyr for data manipulation, and ggplot2 and patchwork for visualization. A colorblind-friendly color palette based on the Okabe-Ito scheme was applied to all scRNA-seq figures

##### Immunohistochemistry

Ovarian and omental tumor sections were stained as previously described. Antigen retrieval was performed by boiling the section in sodium citrate buffer (pH 6.0) for 30 minutes, followed by incubation in 3% hydrogen peroxide at room temperature for 15 minutes to block endogenous peroxide activity; sections were then washed twice in PBS for 5 minutes each and blocked with Background Punisher (BP974, BioCare) for 15 minutes at room temperature. Sections were then incubated at 4°C overnight in primary antibodies for Proliferation cell nuclear antigen (PCNA) (2586, 1:500, Cell Signaling Technology), and CD206 (24595, 1:200, Cell Signaling Technology) and FoxP3 (Cat no. 2653, D6O8R, 1:100, Cell signaling technology) diluted in Da Vinci Green diluent (PD900, BioCare). Detection was performed using the MACH^4^ universal HRP-polymer (M4U534, BioCare) kit or MACH^4^ universal AP-polymer (M4U536, BioCare), followed by the Betazoid DAB Chromogen Kit (BDB2004, BioCare) or Alkaline phosphatase substrate kit (SK-5100, Vectors laboratories) as per the manufacturer’s instructions. Sections were counterstained with hematoxylin for 1 minute, dehydrated through ethanol gradients (70%, 90%, and 100%), and cleared in Xylene for 1 minute, followed by mounting in Paramount TM mounting medium (SP15-100, Fisher Chemicals) for microscopic analysis. A minimum of five regions per tumor core from each mouse were analyzed using QuPath (v0.5.0) Bioimage analysis software ^92^ and presented as a percentage of total cells.

##### Immunofluorescence

For Immunofluorescent detection of immune cells, ovarian and omental sections were processed similarly to IHC up to antigen blocking, omitting peroxidase treatment. Sections were incubated overnight at 4°C with primary antibodies, including Goat anti-mouse CD45 (AF114, 1:100, Novus Biologicals) for immune cells, anti-rabbit Arginase-1 (93668,1:100, Cell Signaling Technology), and rat anti-mouse CD68 (MCA1957, 1:100, Bio-Rad) for macrophage detection For FoxP3 detection, both immunofluorescence and immunohistochemistry approaches were used; immunohistochemistry served as a complementary specificity validation given the technical limitations of multiplex fluorescent staining for nuclear markers on FFPE tissue. Antibodies were diluted in Da Vinci Green diluent. Following primary antibody incubation, sections were washed twice in PBS with 1% tween-20 for 5 minutes and incubated with secondary antibodies, Goat anti-Rabbit IgG Alexa Fluor™ 594 (A11012, 1:500, Invitrogen), Goat anti-Rabbit IgG 488 (A11008, 1:500, Invitrogen), and Donkey anti-rat IgG 488 (A48269TR, 1:500, Invitrogen), for 2 hours at room temperature. Nuclei were counterstained with DAPI for 5 minutes, and sections were mounted using ProLong™ Gold Antifade Mountant (P36930, Thermo Scientific). Multiple Immunofluorescence images (3-5 per tumor core from each mouse) were analyzed using QuPath (v0.5.0) Bioimage analysis software^92^. Double-positive cells were manually counted using Fiji (ImageJ) software by an investigator blinded to the study^93^. Results were presented as the percentage of positive cells per core.

##### Flow cytometry

Ascites fluid was collected from the mice’s peritoneal cavity, and cells were separated from the fluid after centrifugation at 1200 rpm at 4°C. Red blood cells were lysed using RBC lysis with incubation at room temperature for 10 minutes. For the Vismodegib inhibition experiment ovaries and omentum were collected at endpoints (Day 42), after perfusing mice with PBS, digested using Collagenase II (17101015, 1.0 mg/mL, Thermo Fisher) and DNase I (10104159001, 25 μg/mL, Sigma-Aldrich) for 30 minutes at 37°C, passed through 70µm nylon mesh and RBCs were lysed as described above. Freshly isolated cells were stained sequentially with Live/Dead dye (Zombie Yellow^TM^, 423103, Biolegend) for 15 minutes at room temperature followed by Fc-receptor blockade antibody, rat ani-mouse CD16/CD32 (553141, BD Biosciences) for 10 minutes on ice and fluorophore-labeled antibodies at concentrations recommended by manufacturers: Macrophages were identified by gating CD45^+^ cells on F4/80^+^ cells as F4/80^+^CD80^+^ (M1-type) and F4/80^+^CD206^+^ (M2-type) using PerCP/Cyanine5.5 labeled anti-mouse CD45 (103131, Biolegend), FITC labeled anti-mouse F4/80 (123107, Biolegend), APC-labeled anti-mouse CD80 (104714, Biolegend), APC-labeled anti-mouse CD206 (141706, Biolegend). Regulatory T cells were identified by gating CD45^+^ cells on CD3^+^ and CD4^+^ as CD25^+^Foxp3^+^ cells using FITC-labeled anti-mouse CD45 (147709, Biolegend), PerCP 5.5 -labeled anti-mouse CD3 (100218,Biolegend), PE/Cy7-labeled anti-mouse CD4 (100422, Biolegend), APC-labeled anti-mouse CD25 (102012, Biolegend), and PE-labeled anti-mouse Foxp3 (126403, Biolegend).MDSCs were identified by gating CD11b^+^ cells as monocytic (M-MDSC)CD11b^+^Ly6G^-^Ly6C^high^ and polymorphonuclear MDSCs (PMN-MDSC) as CD11b^+^Ly6G^+^Ly6C^low^ using FITC-labeled anti-mouse CD11b(101206, Biolegend), PE-labeled anti-mouse Ly6C (128008, Biolegend), and APC-labeled anti-mouse Ly6G (127614, Biolegend). Cells were incubated with antibodies for 30 minutes on ice in flow cytometry buffer (FACS) containing 2% fetal bovine serum in PBS at the recommended concentrations. For Foxp3 intracellular staining, cells were permeabilized with the True-Nuclear™ Transcription Factor buffer kit (424401, Biolegend) according to the manufacturer’s instructions. Post-staining, cells were resuspended in FACS buffer and analyzed on BD FACSymphony flow cytometer, and data were analyzed using FlowJo^TM^ v10 software (TreeStar Inc., Ashland, OR USA). Comprehensive gating strategies for all flow cytometric analyses are shown in (Supplementary figure 3B, i, vi, xi) and (Supplementary figure 5B, i). Immune cell populations are reported as both percentages (of parent gate) and absolute cell numbers (Supplementary Figures 3B), and per gram of tissue (Supplementary figure 5B) in the case of tissues, calculated using tissue weight and total live-cell counts post-dissociation. We note that absolute count estimates from dissociated tumor tissue are subject to greater technical variability than percentages due to inter-sample differences in enzymatic digestion efficiency, particularly in aged tissues with increased stromal fibrosis.

##### qRT-PCR for Hedgehog pathway inhibition validation

Mesentery tissue from vehicle- and vismodegib-treated mice was collected at endpoint and formalin-fixed and paraffin-embedded following standard histology protocols. RNA was extracted from FFPE sections using RNA isolation kit (Cat. no. 73604, Qiagen) following manufacturer’s instructions, and cDNA was synthesized using reverse transcriptase kit (Cat. no. 95047-500, Quantabio). qRT-PCR was performed using iTaq SYBR Green Super mix (Cat no. 1725125, Bio-Rad) with primers (Supplementary Table C) for mouse Gli1, Ptch1, Hhip, and Rpl13 (housekeeping control). Relative expression was calculated using the 2−ΔΔCt method, normalized to vehicle-treated controls.

##### Bone marrow-derived macrophage (BMDMs) polarization assay

Bone marrow was flushed from femurs and tibias of C57BL/6 female mice. Cells were cultured in DMEM supplemented with 10%FBS and 10ng/mL recombinant murine M-CSF (Cat.no. SRP3221, 10ng/ml, Sigma-Aldrich) for 5 days to differentiate into M0 macrophages. To prepare conditioned media, ID8*Trp*^53^*^−/−^*cells were treated with vismodegib (15 µM) or DMSO vehicle for 48 hours in 1% FBS DMEM; tumor cells were then lysed for qRT-PCR validation of Hedgehog pathway inhibition and the conditioned media was collected, filtered through a 0.22 µm filter, and stored at -80°C until use. M0 BMDMs were treated for 48 hours with: ID8*Trp^53−/−^*conditioned media (50% v/v) from vehicle or conditioned media from vismodegib-pretreated cells (normalized protein content). M-CSF (10ng/ml) was maintained throughout the experiment duration, and IL-4 (Cat. no. 404-ML-025/CF,20ng/ml, R&D systems) was used as a positive control. After 48 hours, total RNA was isolated from BMDMs and qRT-PCR was performed for CD80 and CD206 expression (Mrc1)(Supplementary Table C). Relative expression was calculated using the 2−ΔΔCt method, normalized to the M-CSF-only control.

### QUANTIFICATION AND STATISTICAL ANALYSIS

All data were compared using unpaired Student t-tests and presented as mean ± SEM. One-way ANOVA and two-way ANOVA were performed, followed by Tukey’s and Sidak’s multiple comparison test where applicable. Data was analyzed using Graph Pad Prism 9.0 (La Jolla, CA, USA), and statistical significance was considered at *p* < 0.05. Statistical details for each experiment, including exact n values and significance, are reported in the corresponding figure legends. In all experiments, n represents the number of biologically independent trials (in vitro) or mice (in vivo) unless otherwise stated in the figure legends.

## References

1. Mancebo, G., Sole-Sedeno, J.M., Fabregó, B., Pinto, G., Vizoso, A., Alvarez, M., Sabaté-Garcia, R.A., and Miralpeix, E. (2025). Influence of Age on Treatment and Prognosis in Ovarian Cancer Patients. Cancers (Basel) 17. 10.3390/cancers17091397.

2. Withrow, D.R., Nicholson, B.D., Morris, E.J.A., Wong, M.L., and Pilleron, S. (2023). Age-related differences in cancer relative survival in the United States: A SEER-18 analysis. Int J Cancer 152, 2283–2291. 10.1002/ijc.34463.

3. Labidi-Galy, S.I., Papp, E., Hallberg, D., Niknafs, N., Adleff, V., Noe, M., Bhattacharya, R., Novak, M., Jones, S., Phallen, J., et al. (2017). High grade serous ovarian carcinomas originate in the fallopian tube. Nat Commun 8, 1093. 10.1038/s41467-017-00962-1.

4. Köbel, M., and Kang, E.Y. (2022). The Evolution of Ovarian Carcinoma Subclassification. Cancers (Basel) 14. 10.3390/cancers14020416.

5. Kurman, R.J., and Shih Ie, M. (2010). The origin and pathogenesis of epithelial ovarian cancer: a proposed unifying theory. Am J Surg Pathol 34, 433–443. 10.1097/PAS.0b013e3181cf3d79.

6. Kauff, N.D., Satagopan, J.M., Robson, M.E., Scheuer, L., Hensley, M., Hudis, C.A., Ellis, N.A., Boyd, J., Borgen, P.I., Barakat, R.R., et al. (2002). Risk-reducing salpingo-oophorectomy in women with a BRCA1 or BRCA2 mutation. N Engl J Med 346, 1609–1615. 10.1056/NEJMoa020119.

7. Rebbeck, T.R., Kauff, N.D., and Domchek, S.M. (2009). Meta-analysis of risk reduction estimates associated with risk-reducing salpingo-oophorectomy in BRCA1 or BRCA2 mutation carriers. J Natl Cancer Inst 101, 80–87. 10.1093/jnci/djn442.

8. Domchek, S.M., Friebel, T.M., Singer, C.F., Evans, D.G., Lynch, H.T., Isaacs, C., Garber, J.E., Neuhausen, S.L., Matloff, E., Eeles, R., et al. (2010). Association of risk-reducing surgery in BRCA1 or BRCA2 mutation carriers with cancer risk and mortality. Jama 304, 967–975. 10.1001/jama.2010.1237.

9. Finch, A.P., Lubinski, J., Møller, P., Singer, C.F., Karlan, B., Senter, L., Rosen, B., Maehle, L., Ghadirian, P., Cybulski, C., et al. (2014). Impact of oophorectomy on cancer incidence and mortality in women with a BRCA1 or BRCA2 mutation. J Clin Oncol 32, 1547–1553. 10.1200/jco.2013.53.2820.

10. Ali, A.T., Al-Ani, O., and Al-Ani, F. (2023). Epidemiology and risk factors for ovarian cancer. Prz Menopauzalny 22, 93–104. 10.5114/pm.2023.128661.

11. Briley, S.M., Jasti, S., McCracken, J.M., Hornick, J.E., Fegley, B., Pritchard, M.T., and Duncan, F.E. (2016). Reproductive age-associated fibrosis in the stroma of the mammalian ovary. Reproduction 152, 245–260. 10.1530/rep-16-0129.

12. Maruyama, N., Fukunaga, I., Kogo, T., Endo, T., Fujii, W., Kanai-Azuma, M., Naito, K., and Sugiura, K. (2023). Accumulation of senescent cells in the stroma of aged mouse ovary. J Reprod Dev 69, 328–336. 10.1262/jrd.2023-021.

13. Zhang, Z., Schlamp, F., Huang, L., Clark, H., and Brayboy, L. (2020). Inflammaging is associated with shifted macrophage ontogeny and polarization in the aging mouse ovary. Reproduction 159, 325–337. 10.1530/rep-19-0330.

14. Ben Yaakov, T., Wasserman, T., Aknin, E., and Savir, Y. (2023). Single-cell analysis of the aged ovarian immune system reveals a shift towards adaptive immunity and attenuated cell function. Elife 12. 10.7554/eLife.74915.

15. Loughran, E.A., Leonard, A.K., Hilliard, T.S., Phan, R.C., Yemc, M.G., Harper, E., Sheedy, E., Klymenko, Y., Asem, M., Liu, Y., et al. (2018). Aging Increases Susceptibility to Ovarian Cancer Metastasis in Murine Allograft Models and Alters Immune Composition of Peritoneal Adipose Tissue. Neoplasia 20, 621–631. 10.1016/j.neo.2018.03.007.

16. Hou, X., Zhai, Y., Hu, K., Liu, C.J., Udager, A., Pearce, C.L., Fearon, E.R., and Cho, K.R. (2022). Aging accelerates while multiparity delays tumorigenesis in mouse models of high-grade serous carcinoma. Gynecol Oncol 165, 552–559. 10.1016/j.ygyno.2022.03.030.

17. Flurkey, K., Currer, J.M., and Harrison, D. (2007). Mouse models in aging research. In The mouse in biomedical research, (Elsevier), pp. 637–672.

18. Bhattarai, T., Datta, S., Chaudhuri, P., Bhattacharya, K., and Sengupta, P. (2014). Effect of progesterone supplementation on post-coital unilaterally ovariectomized superovulated mice in relation to implantation and pregnancy. Asian Journal of Pharmaceutical and Clinical Research, 29–31.

19. Wang, S., Lai, X., Deng, Y., and Song, Y. (2020). Correlation between mouse age and human age in anti-tumor research: Significance and method establishment. Life sciences 242, 117242.

20. Hou, X., Zhai, Y., Hu, K., Liu, C.-J., Udager, A., Pearce, C.L., Fearon, E.R., and Cho, K.R. (2022). Aging accelerates while multiparity delays tumorigenesis in mouse models of high-grade serous carcinoma. Gynecologic Oncology 165, 552–559. 10.1016/j.ygyno.2022.03.030.

21. Iyer, S., Zhang, S., Yucel, S., Horn, H., Smith, S.G., Reinhardt, F., Hoefsmit, E., Assatova, B., Casado, J., Meinsohn, M.C., et al. (2021). Genetically Defined Syngeneic Mouse Models of Ovarian Cancer as Tools for the Discovery of Combination Immunotherapy. Cancer Discov 11, 384–407. 10.1158/2159-8290.Cd-20-0818.

22. Integrated genomic analyses of ovarian carcinoma. (2011). Nature 474, 609–615. 10.1038/nature10166.

23. Donovan, M.K.R., D’Antonio-Chronowska, A., D’Antonio, M., and Frazer, K.A. (2020). Cellular deconvolution of GTEx tissues powers discovery of disease and cell-type associated regulatory variants. Nature Communications 11, 955. 10.1038/s41467-020-14561-0.

24. Yuan, X., Zhang, J., Li, D., Mao, Y., Mo, F., Du, W., and Ma, X. (2017). Prognostic significance of tumor-associated macrophages in ovarian cancer: A meta-analysis. Gynecologic oncology 147, 181–187.

25. Hensler, M., Kasikova, L., Fiser, K., Rakova, J., Skapa, P., Laco, J., Lanickova, T., Pecen, L., Truxova, I., and Vosahlikova, S. (2020). M2-like macrophages dictate clinically relevant immunosuppression in metastatic ovarian cancer. Journal for immunotherapy of cancer 8, e000979.

26. Hinshaw, D.C., Hanna, A., Lama-Sherpa, T., Metge, B., Kammerud, S.C., Benavides, G.A., Kumar, A., Alsheikh, H.A., Mota, M., and Chen, D. (2021). Hedgehog signaling regulates metabolism and polarization of mammary tumor-associated macrophages. Cancer research 81, 5425–5437.

27. Roque, D.M., Buza, N., Glasgow, M., Bellone, S., Bortolomai, I., Gasparrini, S., Cocco, E., Ratner, E., Silasi, D.-A., and Azodi, M. (2014). Class III β-tubulin overexpression within the tumor microenvironment is a prognostic biomarker for poor overall survival in ovarian cancer patients treated with neoadjuvant carboplatin/paclitaxel. Clinical & experimental metastasis 31, 101–110.

28. Ma, X. (2020). The omentum, a niche for premetastatic ovarian cancer. Journal of Experimental Medicine 217. 10.1084/jem.20192312.

29. Menjivar, R.E., Nwosu, Z.C., Du, W., Donahue, K.L., Hong, H.S., Espinoza, C., Brown, K., Velez-Delgado, A., Yan, W., and Lima, F. (2023). Arginase 1 is a key driver of immune suppression in pancreatic cancer. Elife 12, e80721.

30. Munder, M. (2009). Arginase: an emerging key player in the mammalian immune system. British journal of pharmacology 158, 638–651.

31. Arlauckas, S.P., Garren, S.B., Garris, C.S., Kohler, R.H., Oh, J., Pittet, M.J., and Weissleder, R. (2018). Arg1 expression defines immunosuppressive subsets of tumor-associated macrophages. Theranostics 8, 5842.

32. Hibbs Jr, J.B., Vavrin, Z., and Taintor, R. (1987). L-arginine is required for expression of the activated macrophage effector mechanism causing selective metabolic inhibition in target cells. The Journal of Immunology 138, 550–565.

33. Colvin, E.K. (2014). Tumor-associated macrophages contribute to tumor progression in ovarian cancer. Frontiers in oncology 4, 137.

34. Carroll, M.J., Kapur, A., Felder, M., Patankar, M.S., and Kreeger, P.K. (2016). M2 macrophages induce ovarian cancer cell proliferation via a heparin binding epidermal growth factor/matrix metalloproteinase 9 intercellular feedback loop. Oncotarget 7, 86608.

35. Recognizing the importance of ovarian aging research. (2022). Nat Aging 2, 1071–1072. 10.1038/s43587-022-00339-0.

36. Hanna, A., Metge, B.J., Bailey, S.K., Chen, D., Chandrashekar, D.S., Varambally, S., Samant, R.S., and Shevde, L.A. (2019). Inhibition of Hedgehog signaling reprograms the dysfunctional immune microenvironment in breast cancer. Oncoimmunology 8, 1548241.

37. Hinshaw, D.C., Hanna, A., Lama-Sherpa, T., Metge, B., Kammerud, S.C., Benavides, G.A., Kumar, A., Alsheikh, H.A., Mota, M., and Chen, D. (2022). Hedgehog signaling regulates metabolism and polarization of mammary tumor-associated macrophages. Cancer Research 82, 2103–2103.

38. Hinshaw, D.C., Benavides, G.A., Metge, B.J., Swain, C.A., Kammerud, S.C., Alsheikh, H.A., Elhamamsy, A., Chen, D., Darley-Usmar, V., Rathmell, J.C., et al. (2023). Hedgehog Signaling Regulates Treg to Th17 Conversion Through Metabolic Rewiring in Breast Cancer. Cancer Immunol Res 11, 687–702. 10.1158/2326-6066.Cir-22-0426.

39. Sabatier, R., Calderon, B., Jr., Lambaudie, E., Chereau, E., Provansal, M., Cappiello, M.A., Viens, P., and Rousseau, F. (2015). Prognostic factors for ovarian epithelial cancer in the elderly: a case-control study. Int J Gynecol Cancer 25, 815–822. 10.1097/igc.0000000000000418.

40. Melamed, A., Bercow, A.S., Bunnell, K., Rauh-Hain, J.A., Wright, J.D., Rice, L.W., and Del Carmen, M.G. (2019). Age-Associated Risk of 90-Day Postoperative Mortality After Cytoreductive Surgery for Advanced Ovarian Cancer. JAMA Surg 154, 669–671. 10.1001/jamasurg.2019.0907.

41. Fourcadier, E., Trétarre, B., Gras-Aygon, C., Ecarnot, F., Daurès, J.P., and Bessaoud, F. (2015). Under-treatment of elderly patients with ovarian cancer: a population based study. BMC Cancer 15, 937. 10.1186/s12885-015-1947-9.

42. Isola, J.V.V., Hense, J.D., Osório, C.A.P., Biswas, S., Alberola-Ila, J., Ocañas, S.R., Schneider, A., and Stout, M.B. (2024). Reproductive Ageing: Inflammation, immune cells, and cellular senescence in the aging ovary. Reproduction 168. 10.1530/rep-23-0499.

43. Cao, L.B., Leung, C.K., Law, P.W.-N., Lv, Y., Ng, C.-H., Liu, H.B., Lu, G., Ma, J.L., and Chan, W.Y. (2020). Systemic changes in a mouse model of VCD-induced premature ovarian failure. Life sciences 262, 118543.

44. Laviolette, L.A., Ethier, J.F., Senterman, M.K., Devine, P.J., and Vanderhyden, B.C. (2011). Induction of a menopausal state alters the growth and histology of ovarian tumors in a mouse model of ovarian cancer. Menopause 18, 549–557. 10.1097/gme.0b013e3181fca1b6.

45. Marion, S.L., Watson, J., Sen, N., Brewer, M.A., Barton, J.K., and Hoyer, P.B. (2013). 7,12-dimethylbenz[a]anthracene-induced malignancies in a mouse model of menopause. Comp Med 63, 6–12.

46. Terry, S., Savagner, P., Ortiz-Cuaran, S., Mahjoubi, L., Saintigny, P., Thiery, J.P., and Chouaib, S. (2017). New insights into the role of EMT in tumor immune escape. Mol Oncol 11, 824–846. 10.1002/1878-0261.12093.

47. Dongre, A., Rashidian, M., Reinhardt, F., Bagnato, A., Keckesova, Z., Ploegh, H.L., and Weinberg, R.A. (2017). Epithelial-to-Mesenchymal Transition Contributes to Immunosuppression in Breast Carcinomas. Cancer Res 77, 3982–3989. 10.1158/0008-5472.Can-16-3292.

48. Dongre, A., and Weinberg, R.A. (2019). New insights into the mechanisms of epithelial-mesenchymal transition and implications for cancer. Nat Rev Mol Cell Biol 20, 69–84. 10.1038/s41580-018-0080-4.

49. Jin, J., Li, Y., Muluh, T.A., Zhi, L., and Zhao, Q. (2021). Identification of CXCL10-Relevant Tumor Microenvironment Characterization and Clinical Outcome in Ovarian Cancer. Front Genet 12, 678747. 10.3389/fgene.2021.678747.

50. Zhou, B., Sun, C., Li, N., Shan, W., Lu, H., Guo, L., Guo, E., Xia, M., Weng, D., Meng, L., et al. (2016). Cisplatin-induced CCL5 secretion from CAFs promotes cisplatin-resistance in ovarian cancer via regulation of the STAT3 and PI3K/Akt signaling pathways. Int J Oncol 48, 2087–2097. 10.3892/ijo.2016.3442.

51. You, Y., Li, Y., Li, M., Lei, M., Wu, M., Qu, Y., Yuan, Y., Chen, T., and Jiang, H. (2018). Ovarian cancer stem cells promote tumour immune privilege and invasion via CCL5 and regulatory T cells. Clin Exp Immunol 191, 60–73. 10.1111/cei.13044.

52. Wertel, I., Surówka, J., Polak, G., Barczyński, B., Bednarek, W., Jakubowicz-Gil, J., Bojarska-Junak, A., and Kotarski, J. (2015). Macrophage-derived chemokine CCL22 and regulatory T cells in ovarian cancer patients. Tumour Biol 36, 4811–4817. 10.1007/s13277-015-3133-8.

53. Marshall, L.A., Marubayashi, S., Jorapur, A., Jacobson, S., Zibinsky, M., Robles, O., Hu, D.X., Jackson, J.J., Pookot, D., Sanchez, J., et al. (2020). Tumors establish resistance to immunotherapy by regulating T(reg) recruitment via CCR4. J Immunother Cancer 8. 10.1136/jitc-2020-000764.

54. Sun, G., and Liu, Y. (2024). Tertiary lymphoid structures in ovarian cancer. Front Immunol 15, 1465516. 10.3389/fimmu.2024.1465516.

55. Dijkgraaf, E.M., Heusinkveld, M., Tummers, B., Vogelpoel, L.T., Goedemans, R., Jha, V., Nortier, J.W., Welters, M.J., Kroep, J.R., and van der Burg, S.H. (2013). Chemotherapy alters monocyte differentiation to favor generation of cancer-supporting M2 macrophages in the tumor microenvironment. Cancer Res 73, 2480–2492. 10.1158/0008-5472.Can-12-3542.

56. Bellora, F., Castriconi, R., Dondero, A., Pessino, A., Nencioni, A., Liggieri, G., Moretta, L., Mantovani, A., Moretta, A., and Bottino, C. (2014). TLR activation of tumor-associated macrophages from ovarian cancer patients triggers cytolytic activity of NK cells. Eur J Immunol 44, 1814–1822. 10.1002/eji.201344130.

57. Isola, J.V.V., Ocañas, S.R., Hubbart, C.R., Ko, S., Mondal, S.A., Hense, J.D., Carter, H.N.C., Schneider, A., Kovats, S., Alberola-Ila, J., et al. (2024). A single-cell atlas of the aging mouse ovary. Nature Aging 4, 145–162. 10.1038/s43587-023-00552-5.

58. Curiel, T.J., Coukos, G., Zou, L., Alvarez, X., Cheng, P., Mottram, P., Evdemon-Hogan, M., Conejo-Garcia, J.R., Zhang, L., Burow, M., et al. (2004). Specific recruitment of regulatory T cells in ovarian carcinoma fosters immune privilege and predicts reduced survival. Nature Medicine 10, 942–949. 10.1038/nm1093.

59. Condamine, T., Ramachandran, I., Youn, J.I., and Gabrilovich, D.I. (2015). Regulation of tumor metastasis by myeloid-derived suppressor cells. Annu Rev Med 66, 97–110. 10.1146/annurev-med-051013-052304.

60. Zou, G., Wang, J., Xu, X., Xu, P., Zhu, L., Yu, Q., Peng, Y., Guo, X., Li, T., and Zhang, X. (2021). Cell subtypes and immune dysfunction in peritoneal fluid of endometriosis revealed by single-cell RNA-sequencing. Cell Biosci 11, 98. 10.1186/s13578-021-00613-5.

61. Kubicka, U., Olszewski, W.L., Tarnowski, W., Bielecki, K., Ziółkowska, A., and Wierzbicki, Z. (1996). Normal human immune peritoneal cells: subpopulations and functional characteristics. Scand J Immunol 44, 157–163. 10.1046/j.1365-3083.1996.d01-297.x.

62. Yona, S., Kim, K.W., Wolf, Y., Mildner, A., Varol, D., Breker, M., Strauss-Ayali, D., Viukov, S., Guilliams, M., Misharin, A., et al. (2013). Fate mapping reveals origins and dynamics of monocytes and tissue macrophages under homeostasis. Immunity 38, 79–91. 10.1016/j.immuni.2012.12.001.

63. Louwe, P.A., Badiola Gomez, L., Webster, H., Perona-Wright, G., Bain, C.C., Forbes, S.J., and Jenkins, S.J. (2021). Recruited macrophages that colonize the post-inflammatory peritoneal niche convert into functionally divergent resident cells. Nat Commun 12, 1770. 10.1038/s41467-021-21778-0.

64. Zhang, Z., Huang, L., and Brayboy, L. (2021). Macrophages: an indispensable piece of ovarian health. Biol Reprod 104, 527–538. 10.1093/biolre/ioaa219.

65. Gampala, S., and Yang, J.Y. (2021). Hedgehog Pathway Inhibitors against Tumor Microenvironment. Cells 10. 10.3390/cells10113135.

66. Jing, J., Wu, Z., Wang, J., Luo, G., Lin, H., Fan, Y., and Zhou, C. (2023). Hedgehog signaling in tissue homeostasis, cancers, and targeted therapies. Signal Transduct Target Ther 8, 315. 10.1038/s41392-023-01559-5.

67. Bhadricha, H., Patel, V., Singh, A.K., Savardekar, L., Patil, A., Surve, S., and Desai, M. (2021). Increased frequency of Th17 cells and IL-17 levels are associated with low bone mineral density in postmenopausal women. Sci Rep 11, 16155. 10.1038/s41598-021-95640-0.

68. Kapeni, C., O’Brien, L., Sabirova, D., Cast, O., Carbonaro, V., Clark-Leonard, S., Machel, A.C., Beke, F., McDonald, S., Fife, K., and de la Roche, M. (2025). Noncanonical Hedgehog signaling through Smoothened controls cytotoxic T cell migration in the tumor microenvironment. Science Immunology 10, eadr3127. doi:10.1126/sciimmunol.adr3127.

69. Kaye, S.B., Fehrenbacher, L., Holloway, R., Amit, A., Karlan, B., Slomovitz, B., Sabbatini, P., Fu, L., Yauch, R.L., Chang, I., and Reddy, J.C. (2012). A phase II, randomized, placebo-controlled study of vismodegib as maintenance therapy in patients with ovarian cancer in second or third complete remission. Clin Cancer Res 18, 6509–6518. 10.1158/1078-0432.Ccr-12-1796.

70. Evans, E.T., Page, E.F., Choi, A.S., Shonibare, Z., Kahn, A.G., Arend, R.C., and Mythreye, K. (2024). Activin levels correlate with lymphocytic infiltration in epithelial ovarian cancer. Cancer Med 13, e7368. 10.1002/cam4.7368.

71. Luo, X., Fu, Y., Loza, A.J., Murali, B., Leahy, K.M., Ruhland, M.K., Gang, M., Su, X., Zamani, A., Shi, Y., et al. (2016). Stromal-Initiated Changes in the Bone Promote Metastatic Niche Development. Cell Rep 14, 82–92. 10.1016/j.celrep.2015.12.016.

72. Mayer, L.P., Devine, P.J., Dyer, C.A., and Hoyer, P.B. (2004). The follicle-deplete mouse ovary produces androgen. Biol Reprod 71, 130–138. 10.1095/biolreprod.103.016113.

73. Wong, H., Alicke, B., West, K.A., Pacheco, P., La, H., Januario, T., Yauch, R.L., de Sauvage, F.J., and Gould, S.E. (2011). Pharmacokinetic-pharmacodynamic analysis of vismodegib in preclinical models of mutational and ligand-dependent Hedgehog pathway activation. Clin Cancer Res 17, 4682–4692. 10.1158/1078-0432.Ccr-11-0975.

74. Carreno, G., Boult, J.K.R., Apps, J., Gonzalez-Meljem, J.M., Haston, S., Guiho, R., Stache, C., Danielson, L.S., Koers, A., Smith, L.M., et al. (2019). SHH pathway inhibition is protumourigenic in adamantinomatous craniopharyngioma. Endocr Relat Cancer 26, 355–366. 10.1530/erc-18-0538.

75. Vichai, V., and Kirtikara, K. (2006). Sulforhodamine B colorimetric assay for cytotoxicity screening. Nat Protoc 1, 1112–1116. 10.1038/nprot.2006.179.

76. Subramanian, A., Tamayo, P., Mootha, V.K., Mukherjee, S., Ebert, B.L., Gillette, M.A., Paulovich, A., Pomeroy, S.L., Golub, T.R., Lander, E.S., and Mesirov, J.P. (2005). Gene set enrichment analysis: A knowledge-based approach for interpreting genome-wide expression profiles. Proceedings of the National Academy of Sciences 102, 15545–15550. doi:10.1073/pnas.0506580102.

77. Koopmans, F., van Nierop, P., Andres-Alonso, M., Byrnes, A., Cijsouw, T., Coba, M.P., Cornelisse, L.N., Farrell, R.J., Goldschmidt, H.L., Howrigan, D.P., et al. (2019). SynGO: An Evidence-Based, Expert-Curated Knowledge Base for the Synapse. Neuron 103, 217–234.e214. 10.1016/j.neuron.2019.05.002.

78. Li, T., Fu, J., Zeng, Z., Cohen, D., Li, J., Chen, Q., Li, B., and Liu, X.S. (2020). TIMER2.0 for analysis of tumor-infiltrating immune cells. Nucleic Acids Res 48, W509–w514. 10.1093/nar/gkaa407.

79. Kernick, K., Woudstra, R., Berjanskii, M., MacKay, S., and Wishart, D.S. (2025). Heatmapper2: web-enabled heat mapping made easy. Nucleic Acids Res 53, W316–w323. 10.1093/nar/gkaf385.

80. Tang, D., Chen, M., Huang, X., Zhang, G., Zeng, L., Zhang, G., Wu, S., and Wang, Y. (2023). SRplot: A free online platform for data visualization and graphing. PLoS One 18, e0294236. 10.1371/journal.pone.0294236.

81. Gao, J., Aksoy, B.A., Dogrusoz, U., Dresdner, G., Gross, B., Sumer, S.O., Sun, Y., Jacobsen, A., Sinha, R., Larsson, E., et al. (2013). Integrative analysis of complex cancer genomics and clinical profiles using the cBioPortal. Sci Signal 6, pl1. 10.1126/scisignal.2004088.

82. de Bruijn, I., Kundra, R., Mastrogiacomo, B., Tran, T.N., Sikina, L., Mazor, T., Li, X., Ochoa, A., Zhao, G., Lai, B., et al. (2023). Analysis and Visualization of Longitudinal Genomic and Clinical Data from the AACR Project GENIE Biopharma Collaborative in cBioPortal. Cancer Res 83, 3861–3867. 10.1158/0008-5472.Can-23-0816.

83. Cerami, E., Gao, J., Dogrusoz, U., Gross, B.E., Sumer, S.O., Aksoy, B.A., Jacobsen, A., Byrne, C.J., Heuer, M.L., Larsson, E., et al. (2012). The cBio cancer genomics portal: an open platform for exploring multidimensional cancer genomics data. Cancer Discov 2, 401–404. 10.1158/2159-8290.Cd-12-0095.

84. Stuart, T., Butler, A., Hoffman, P., Hafemeister, C., Papalexi, E., Mauck, W.M., 3rd, Hao, Y., Stoeckius, M., Smibert, P., and Satija, R. (2019). Comprehensive Integration of Single-Cell Data. Cell 177, 1888–1902.e1821. 10.1016/j.cell.2019.05.031.

85. Rooney, M.S., Shukla, S.A., Wu, C.J., Getz, G., and Hacohen, N. (2015). Molecular and genetic properties of tumors associated with local immune cytolytic activity. Cell 160, 48–61. 10.1016/j.cell.2014.12.033.

86. Ayers, M., Lunceford, J., Nebozhyn, M., Murphy, E., Loboda, A., Kaufman, D.R., Albright, A., Cheng, J.D., Kang, S.P., Shankaran, V., et al. (2017). IFN-γ-related mRNA profile predicts clinical response to PD-1 blockade. J Clin Invest 127, 2930–2940. 10.1172/jci91190.

87. Auslander, N., Zhang, G., Lee, J.S., Frederick, D.T., Miao, B., Moll, T., Tian, T., Wei, Z., Madan, S., Sullivan, R.J., et al. (2018). Robust prediction of response to immune checkpoint blockade therapy in metastatic melanoma. Nat Med 24, 1545–1549. 10.1038/s41591-018-0157-9.

88. Spranger, S., Bao, R., and Gajewski, T.F. (2015). Melanoma-intrinsic β-catenin signalling prevents anti-tumour immunity. Nature 523, 231–235. 10.1038/nature14404.

89. Hugo, W., Zaretsky, J.M., Sun, L., Song, C., Moreno, B.H., Hu-Lieskovan, S., Berent-Maoz, B., Pang, J., Chmielowski, B., Cherry, G., et al. (2016). Genomic and Transcriptomic Features of Response to Anti-PD-1 Therapy in Metastatic Melanoma. Cell 165, 35–44. 10.1016/j.cell.2016.02.065.

90. Cabrita, R., Lauss, M., Sanna, A., Donia, M., Skaarup Larsen, M., Mitra, S., Johansson, I., Phung, B., Harbst, K., Vallon-Christersson, J., et al. (2020). Tertiary lymphoid structures improve immunotherapy and survival in melanoma. Nature 577, 561–565. 10.1038/s41586-019-1914-8.

91. Peng, J., Sun, B.F., Chen, C.Y., Zhou, J.Y., Chen, Y.S., Chen, H., Liu, L., Huang, D., Jiang, J., Cui, G.S., et al. (2019). Single-cell RNA-seq highlights intra-tumoral heterogeneity and malignant progression in pancreatic ductal adenocarcinoma. Cell Res 29, 725–738. 10.1038/s41422-019-0195-y.

92. Bankhead, P., Loughrey, M.B., Fernández, J.A., Dombrowski, Y., McArt, D.G., Dunne, P.D., McQuaid, S., Gray, R.T., Murray, L.J., Coleman, H.G., et al. (2017). QuPath: Open source software for digital pathology image analysis. Scientific Reports 7, 16878. 10.1038/s41598-017-17204-5.

93. Schindelin, J., Arganda-Carreras, I., Frise, E., Kaynig, V., Longair, M., Pietzsch, T., Preibisch, S., Rueden, C., Saalfeld, S., Schmid, B., et al. (2012). Fiji: an open-source platform for biological-image analysis. Nature Methods 9, 676–682. 10.1038/nmeth.2019.

