## Supplementary text and figures for "Aging increases ovarian cancer growth, metastasis, and immunosuppression that can be alleviated by inhibiting hedgehog signaling"

**SUPPLEMENTARY FIGURE 1.A**

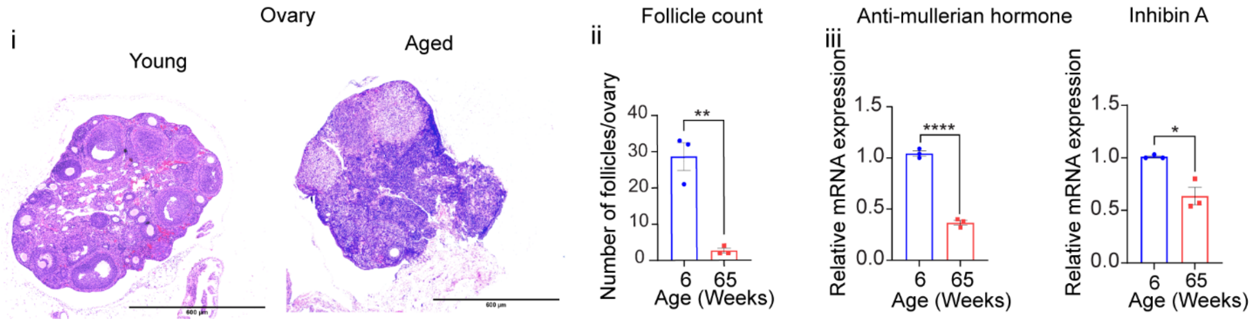


**Supplementary Figure 1. (A)**(i-ii) Representative images of histological analysis (H&E staining) of ovaries from aged (65-week-old) and young (6-week-old) mice, along with follicle count analysis. (iii) Semi-quantitative PCR showing *AMH* and *INHA* (Inhibin A) and mRNA expression levels in ovaries from aged and young mice. All data are mean ± SEM (n=3). *p<0.05, **p<0.01, ***p<0.001, ****p<0.0001, unpaired t test.

**SUPPLEMENTARY FIGURE 1.B**

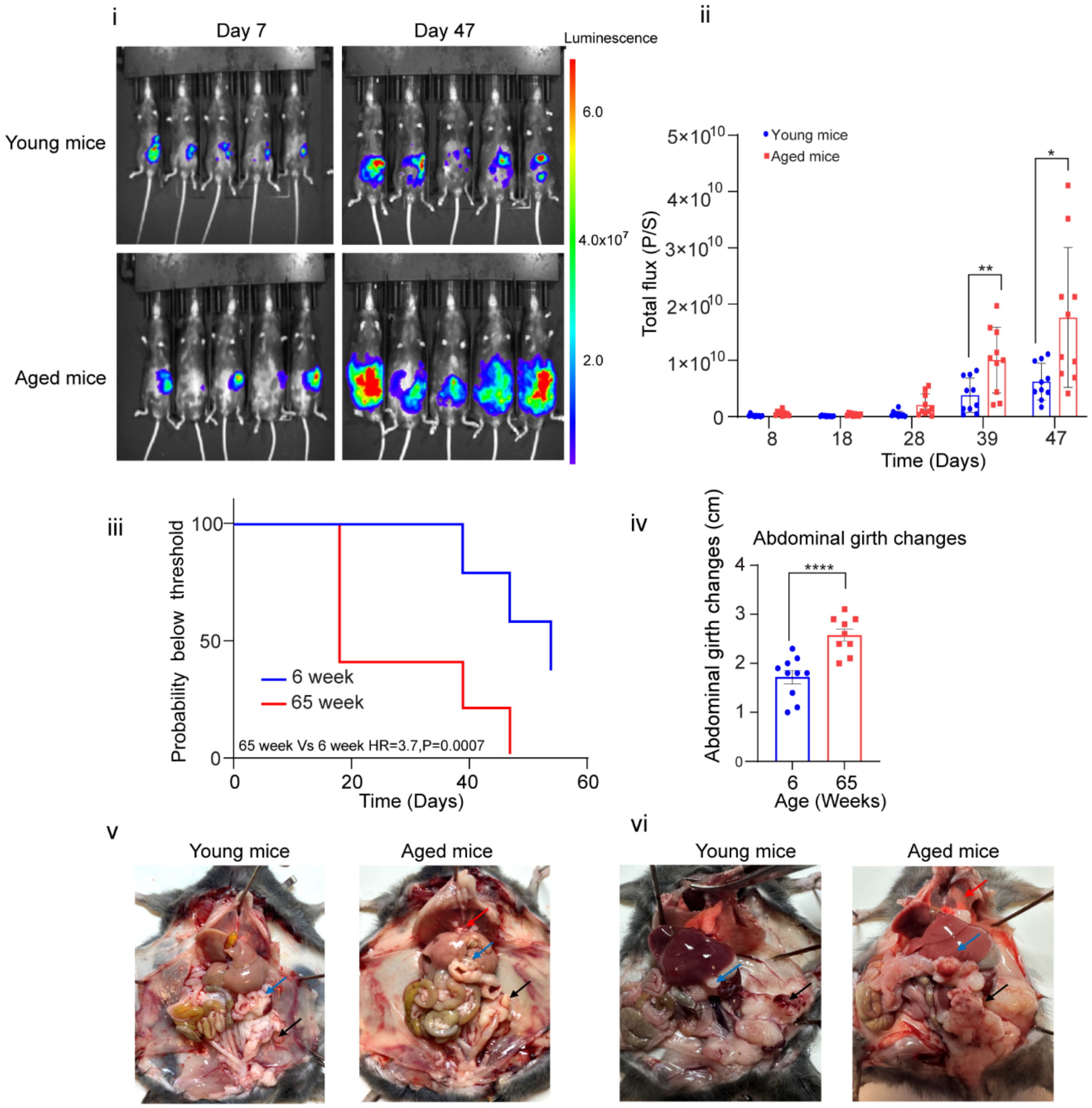


**Supplementary Figure 1 (B)**(i-ii) Representative images at day 39 and day 47 and quantification of tumor growth kinetics from bioluminescence imaging (BLI) from day 7 to day 47 following intrabursal implantation of ID8*Trp53* $^{-/-}$ cells into mouse ovaries in either aged (65 week old) or young mouse hosts (6 week old). (iii) Probability plot based on metastatic tumor area in the peritoneum showing probability of remaining below the threshold of metastatic tumor area in young mice from day 7 to day 54 post-intrabursal tumor cell implantation. (iv) Abdominal girth measurements at endpoint. (v) Representative images of young and aged mice at endpoint ( day 55) post ID8*Trp53* $^{-/-}$ cell implantation into ovaries showing visible tumor growth (vi) Representative images of tumor bearing young and aged mice at endpoint (day 42) following intrabursal implantation of murine fallopian tube-derived PPNM cells . Red arrows indicate diaphragm tumors, blue arrows indicate omental tumors, and black arrows indicate ovarian tumors. All data are mean ± SEM (n=10); *p<0.05, **p<0.01, ***p<0.001, ****p<0.0001, unpaired t test.

**SUPPLEMENTARY FIGURE 1.C**

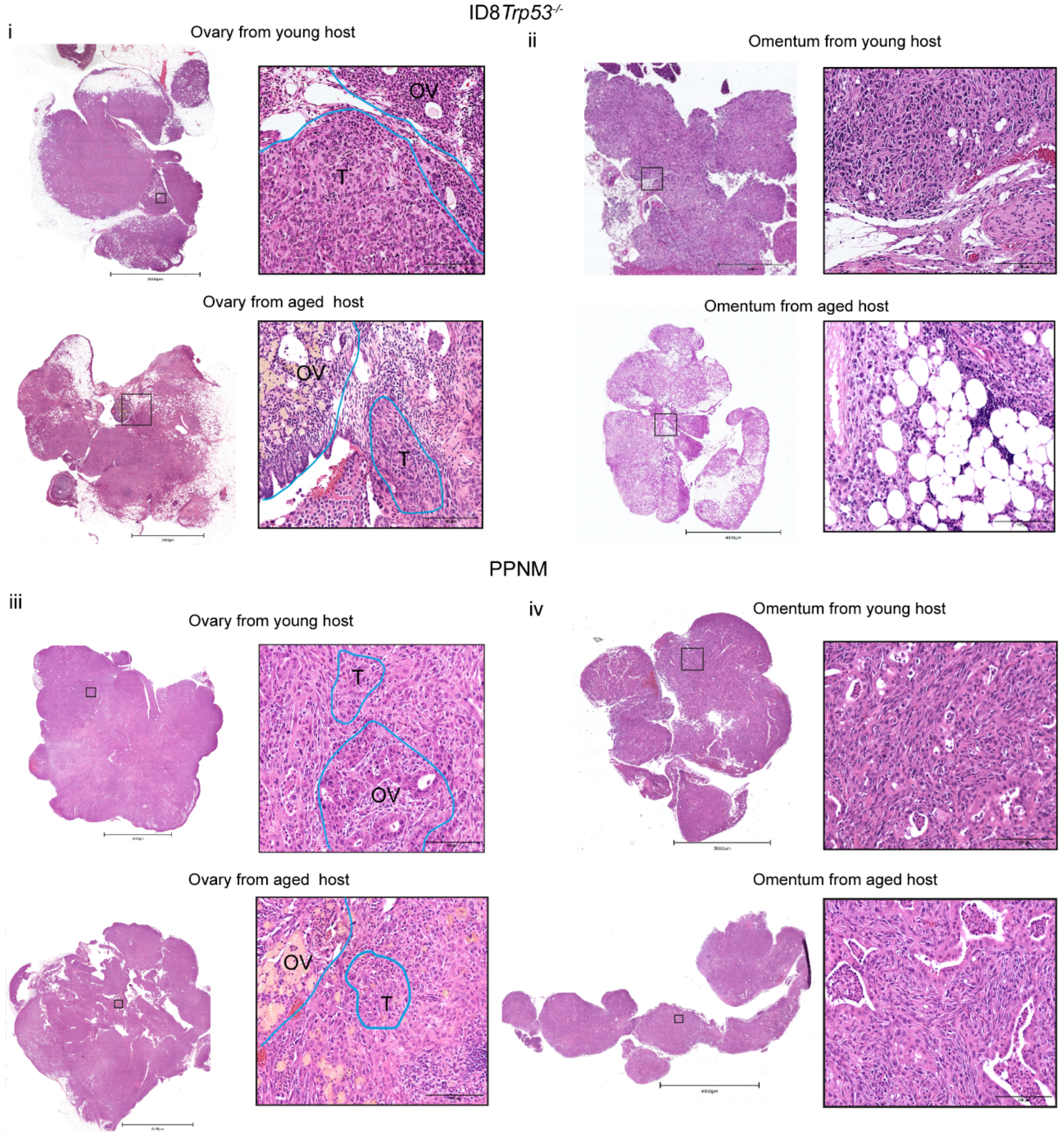


**Supplementary Figure 1. (C)**(i-iv) Ovarian and omental tumors from young and aged hosts intrabursally implanted with ID8*Trp53*$^{-/-}$ or PPNM cells**.** The ovaries (OV) and tumor cells (T) are highlighted.

**SUPPLEMENTARY FIGURE 1.D**

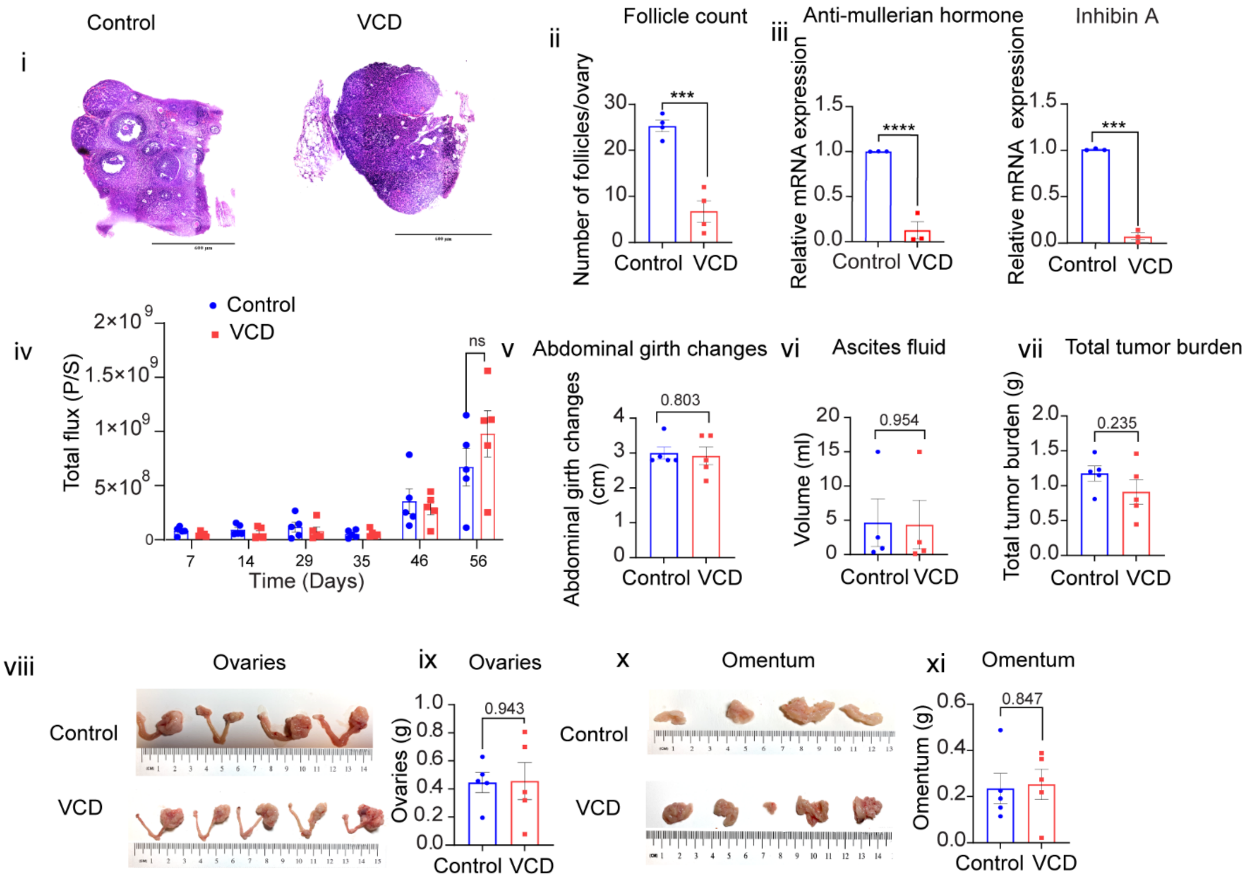


**Supplementary Figure 1.(D)**(i-ii) Histological examination (H&E staining) of ovaries from Vinylcyclohexene dioxide (VCD)-treated and control mice showing marked follicle depletion in the VCD group.(iii) Semi-quantitative PCR analysis of *AMH* and *INHA* mRNA expression in VCD-treated and control ovaries (n=3). (iv) Whole-body bioluminescence imaging (BLI) quantification of tumor growth in 6-week-old C57BL/6 female mice following intrabursal implantation of ID8*Trp53*$^{-/-}$cells, monitored up to day 56. (v) Abdominal girth measurements (vi) quantification of ascites fluid volume (p=0.954) and (vii) Total tumor burden are presented. (viii-xi) Representative images and quantification of ovaries and omentum at endpoint day 67. All data are presented as mean±SEM (n=5) *p<0.05, **p<0.01, ***p<0.001, ****p<0.0001, unpaired t test.

**SUPPLEMENTARY FIGURE 1.E**

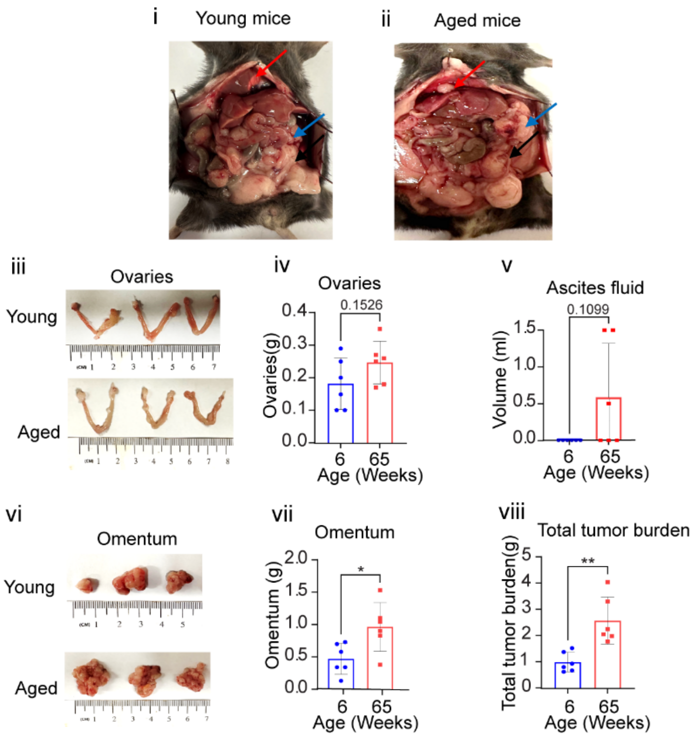


**Supplementary Figure 1. (E)**(i-ii) Representative images of young and aged mice at endpoint (day 30) following intraperitoneal implantation of murine fallopian tube-derived PPNM cells showing tumor formation in multiple sites. Red arrows indicate diaphragm tumors, blue arrows indicate omental tumors, and black arrows indicate ovarian tumors.(iii)&(vi) Representative images of tumor burden in the ovarian and omentum of the mice of indicated age groups (iv) Quantifications of ovary weights (v) ascites fluid (vii) omental weights (viii) total tumor burden on day 30 post intraperitoneal implantation of PPNM cells in young and aged mice (n=6). All data are mean ± SEM (n=6); *$p<0.05$, **$p<0.01$, ***$p<0.001$, ****$p<0.0001$, unpaired t test.

SUPPLEMENTARY FIGURE 2.A

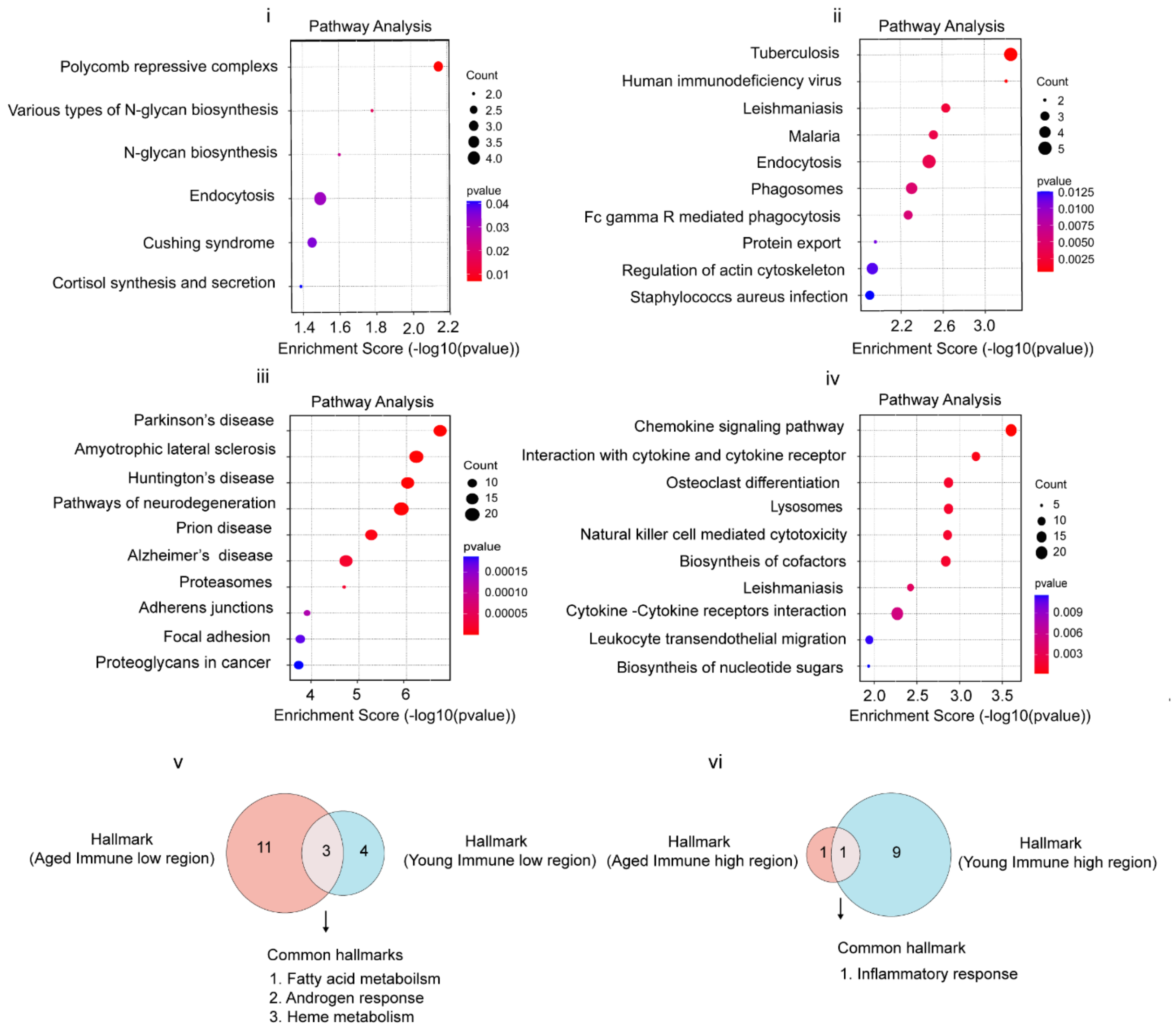


**Supplementary Figure 2. (A)** KEGG pathway in (i) CD45$^{+}$enriched (ii) CD45$^{+}$depleted regions from young host ovarian tumors and KEGG pathways in (iii) CD45$^{+}$-enriched (iv) CD45$^{+}$-depleted regions from aged host ovarian tumors. (v-vi) Common and unique hallmark pathways between aged or young immune regions : (v) low (vi) high regions.

**SUPPLEMENTARY FIGURE 2.B**

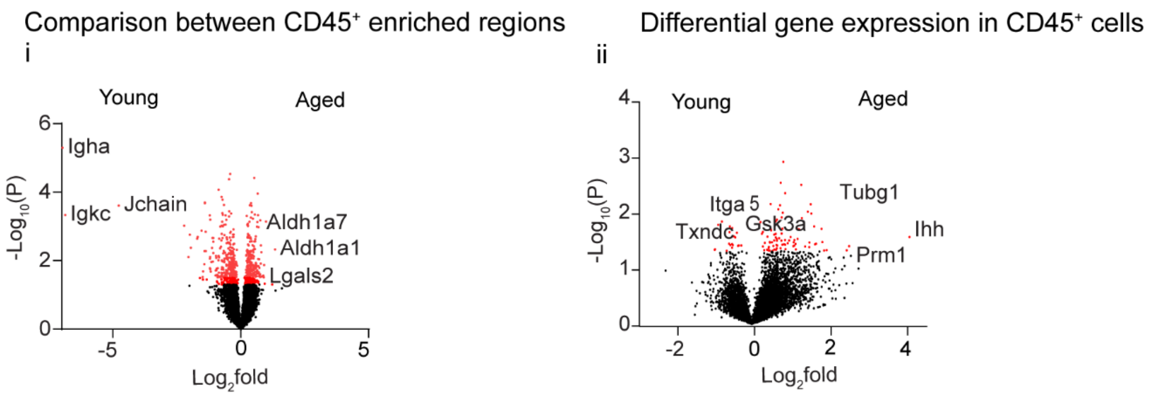


**Supplementary Figure 2.(B)** Volcano plot showing **(i)** Differentially expressed genes in CD45$^+$ enriched regions of ovarian tumors of young or aged host. **(ii)** Differential gene expression in the CD45$^+$ cells from young vs aged host ovarian tumors ($p<0.05$).

### SUPPLEMENTARY FIGURE 3.A

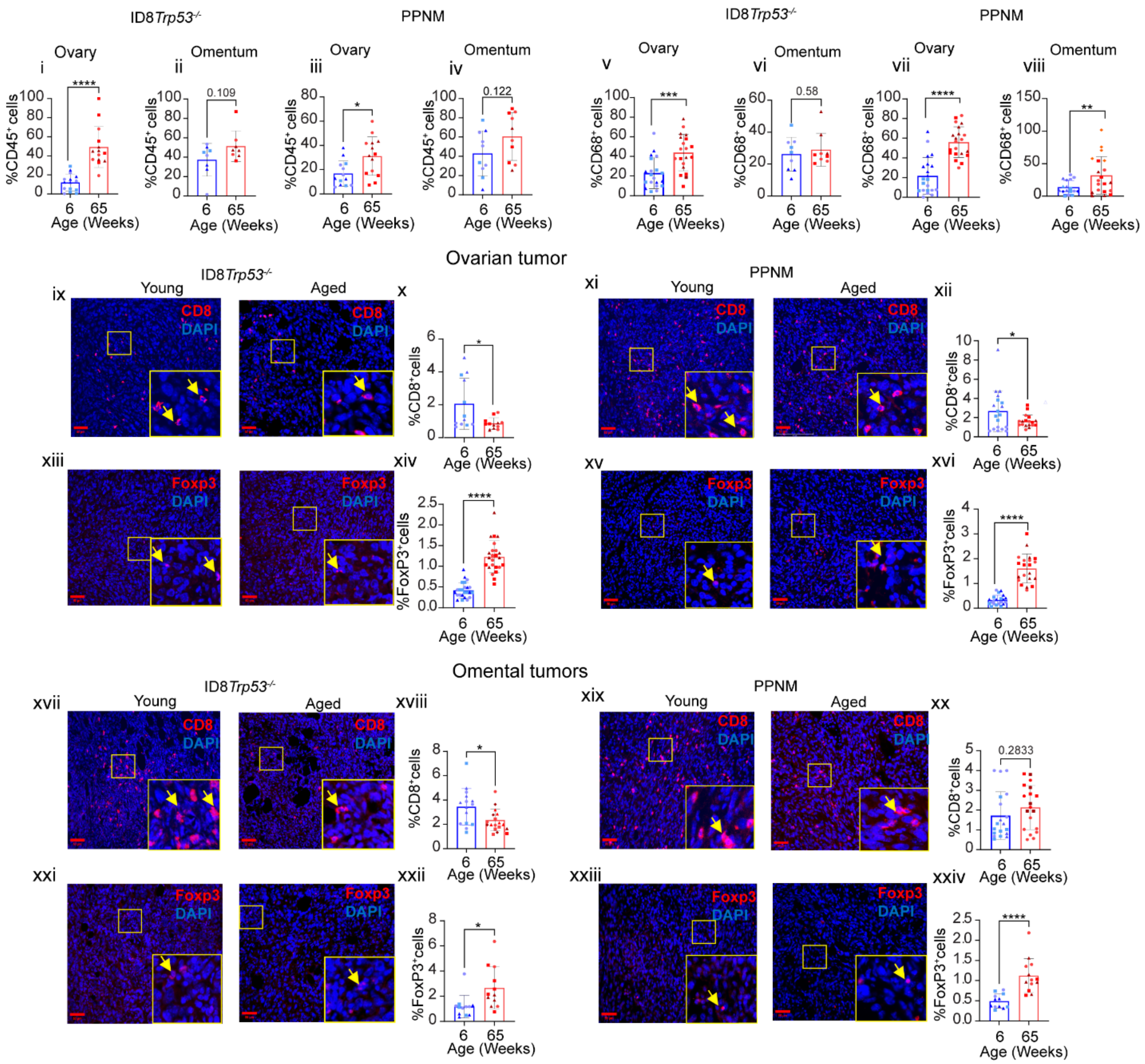


**Supplementary Figure 3. (A i-viii)** Quantification of percentage CD45$^{+}$ cells from ID8*Trp53*$^{-/-}$ or PPNM ovarian or omental tumors as indicated (n=3). **(ix-xii)** Representative immunofluorescence images from ovarian ID8*Trp53*$^{-/-}$ or PPNM tumors showing CD8$^{+}$ (red) and DAPI (blue). Adjacent graphs show quantification of percentage cytotoxic CD8$^{+}$ cells in both models. **(xiii-xvi)** Representative immunofluorescence images of FoxP3$^{+}$(red) and DAPI (blue) from ovarian ID8*Trp53*$^{-/-}$ or PPNM tumors. Adjacent graphs show quantification of percentage FoxP3$^{+}$ cells in both models. **(xvii-xx)** Representative immunofluorescence images from omental

ID8*Trp53*$^{-/-}$ or PPNM tumors showing $CD8^+$(red) and DAPI (Blue). Adjacent graphs show quantification of percentage cytotoxic $CD8^+$ cells in both models. **(xxi-xxiv)** Representative immunofluorescence images of $FoxP3^+$(red) and DAPI (blue) from omental ID8*Trp53*$^{-/-}$ or PPNM tumors. Adjacent graphs show quantification of percentage $FoxP3^+$ cells in both models. Color codes and symbols indicate the number of images analyzed per mouse in each group. (n=3). All data are mean ± SEM, *p<0.05, **p<0.01, ***p<0.001, ****p<0.0001, unpaired t test.

### SUPPLEMENTARY FIGURE 3.B

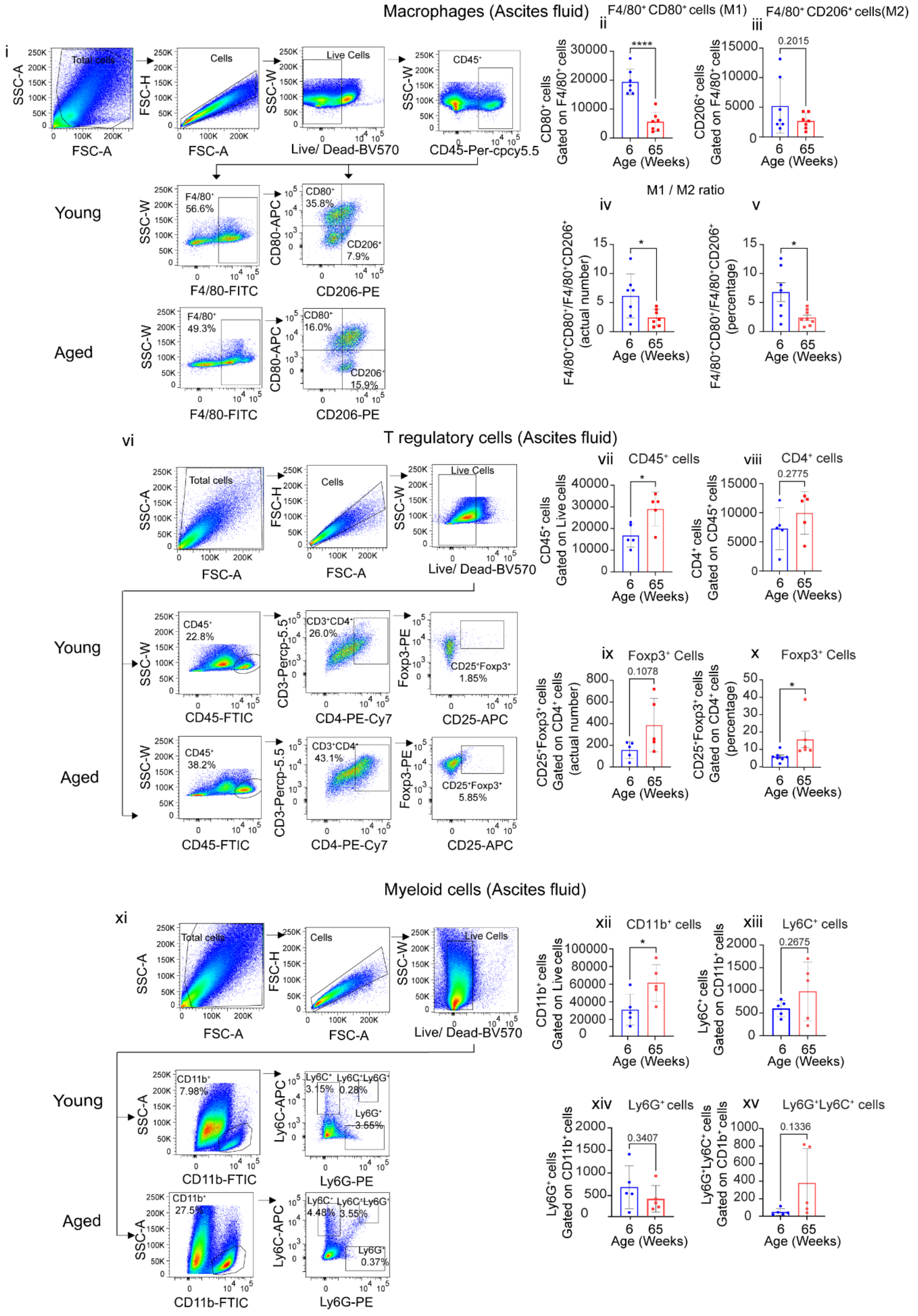

**Supplementary Figure 3. (B)** (i) Gating strategy of F4/80$^+$CD206$^+$ and F4/80$^+$CD80$^+$ cells over CD45$^+$ cells from ascites, (ii) Actual number of F4/80$^+$CD80$^+$ (M1 macrophages) cells gated on F4/80$^+$ cells from ascites. (iii) Actual number of F4/80$^+$CD206$^+$ (M2 macrophages) cells gated on F4/80$^+$ cells. (iv) Ratio of the actual number of F4/80$^+$CD80$^+$(M1)/F4/80$^+$CD206$^+$(M2) cells from ascites. (v) Percentage ratio of F4/80$^+$CD80$^+$(M1)/F4/80$^+$CD206$^+$(M2) cells (n=6). **(**vi) Gating strategy of CD25$^+$ and Foxp3$^+$ cells on CD45$^+$ and CD3$^+$CD4$^+$ cells from ascites. (vii) Actual number of CD45$^+$ cells gated on live cells from ascites. (viii) Actual number of CD4$^+$ cells gated on CD45$^+$ cells from ascites. (ix) Actual number of CD4$^+$CD25$^+$Foxp3$^+$ gated on CD4$^+$ cells . (x) Percentage of CD4$^+$CD25$^+$FoxP3$^+$ gated on CD4$^+$ cells from ascites (n=5). (xi) Gating strategy of MDSCs cells. (xii) Actual number of CD11b$^+$ gated on live cells . (xiii) Actual number of M-MDSCs as CD11b$^+$Ly6G$^-$Ly6C$^{high}$ cell gated on CD11b$^+$ cells. (xiv) Actual number of PMN:MDSCs as CD11b$^+$Ly6G$^+$Ly6C$^{low}$ cells gated on CD11b$^+$ cells from ascites. (xv) Actual number of CD11b$^+$Ly6$^+$G Ly6C$^+$ double positive cells from ascites (n=5). All data are mean ± SEM, *p<0.05, **p<0.01, ***p<0.001, ***p<0.0001, unpaired t-test**.**

**SUPPLEMENTARY FIGURE 3.C**

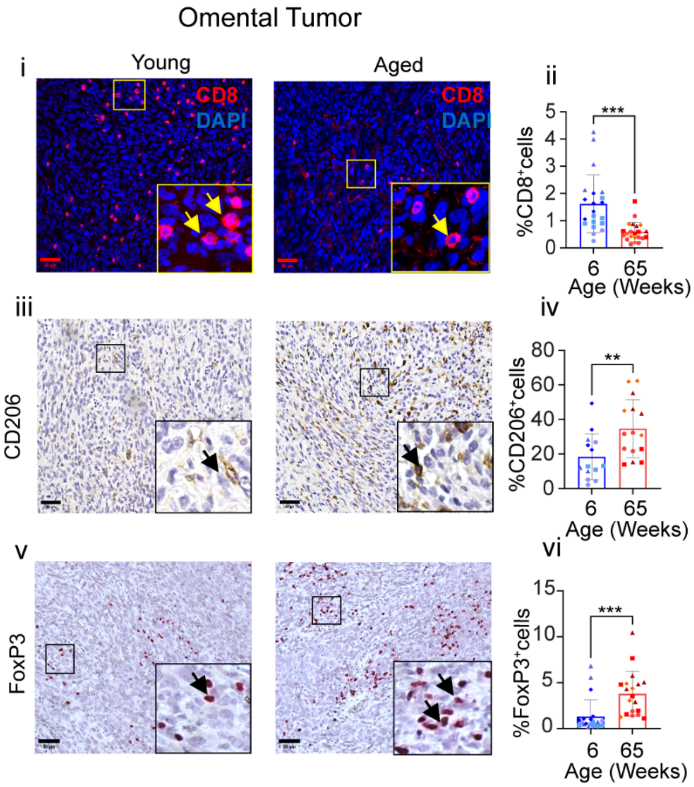


**Supplementary Figure 3. (C)**(i) Representative immunofluorescence images and (ii) quantifications of CD8+ cells in the omentum from intraperitoneally (i.p) implanted PPNM cells (iii) representative immunohistochemistry images of CD206+ cells and (iv) quantifications in omentum from i.p implanted PPNM cells (v) representative FoxP3 immunohistochemistry images and (vi) quantifications in omentum from intraperitoneally (i.p) implanted PPNM cells. (n=4), all data are mean ± SEM, *p<0.05, **p<0.01, ***p<0.001, ***p<0.0001, unpaired t-test.

**SUPPLEMENTARY FIGURE 4**

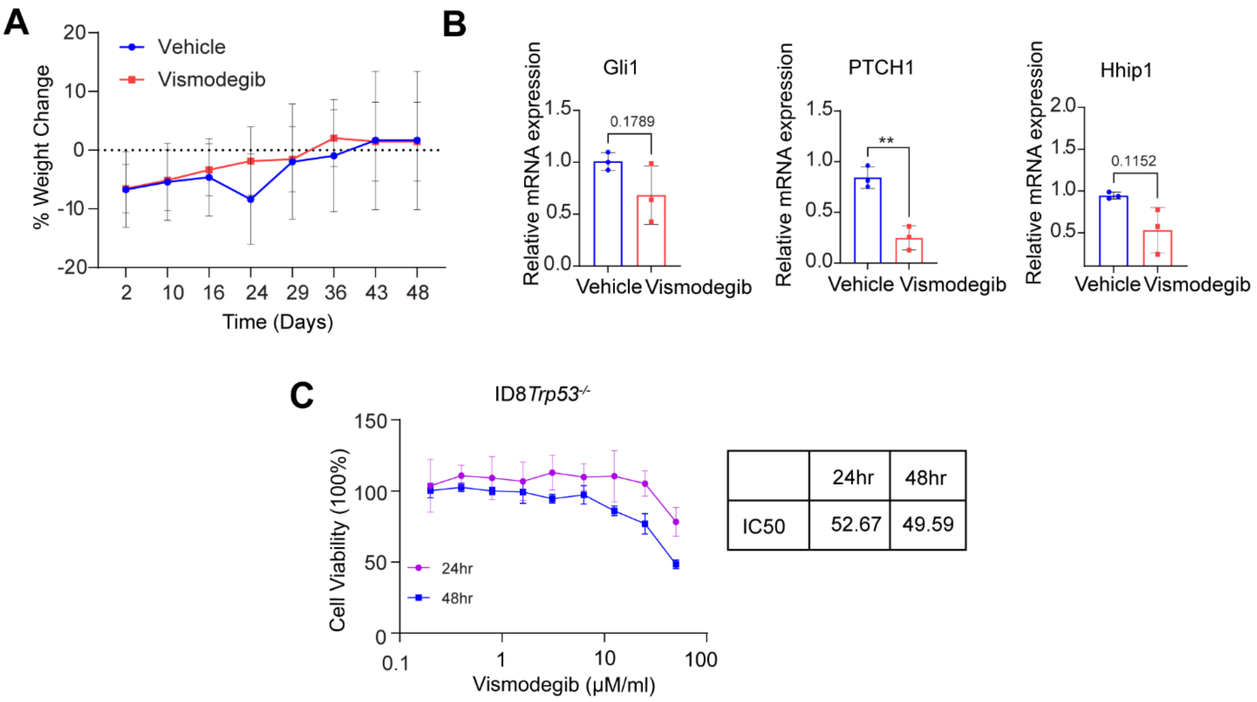


**Supplementary Figure 4. (A)** Percent Weight change from day 2 post ID8*Trp53*$^{-/-}$ cell implantation to day 48 in vehicle and Vismodegib treated mice. **(B)** mRNA quantification of hedgehog pathway genes *Gli1, Ptch1,* and *Hhip1* in mesentery tissues from vismodegib and vehicle-treated mice. Two-day dose response curve of ID8*Trp53*$^{-/-}$cells treated with increasing concentration of vismodegib (0-50µM/ml). Cell viability data are expressed as percentage SRB measurements relative to vehicle-treated cells. IC50 values from three independent experiments with six replicates per drug concentration are shown in the table from either 24hr or 48hr of drug treatment.

### SUPPLEMENTARY FIGURE 5

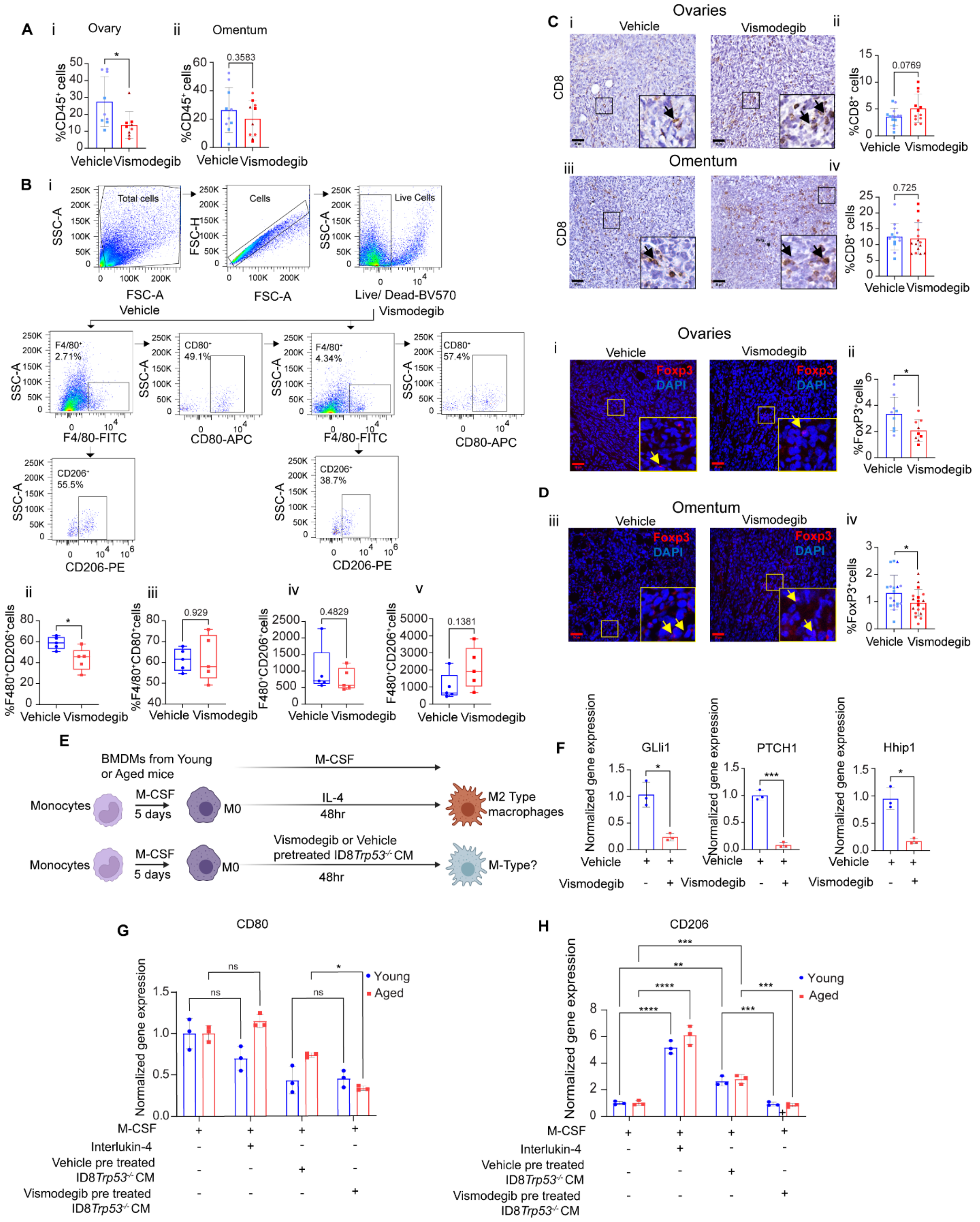

**Supplementary Figure 5. (A)** Quantification of percent $CD45^+$ cells in **(i)** ovary and **(ii)** omental tumors of vismodegib or vehicle-treated mice. **(B)**(i) Flow cytometry gating strategy for macrophages population, $F4/80^+$ (M0) ,$CD80^+$ (M1) and $CD206^+$ (M2) cells over live cells**.** Percentage of (ii) $F4/80^+CD206^+$ or **(**iii) $F4/80^+CD80^+$cells in ovaries and (iv) Actual number of $F4/80^+CD206^+$ and (v) $F4/80^+CD80^+$cells per gram of ovarian tissue. **(C i-iv)** Representative $CD8^+$ immunohistochemistry images in ovaries and omental tumors as indicated with adjacent quantification. **(D)** (i-iv) Representative $FoxP3^+$ (red) and DAPI (Blue) immunofluorescence images cells in ovarian and omental tumors with adjacent graph showing quantification of percentage $FoxP3^+$ cells. Each color and pattern indicates the core analyzed from the mouse in each group (n=3). All data are mean ± SEM (n=5): *p<0.05, **p<0.01, ***p<0.001, ****p<0.0001, unpaired t test. **(E)** Schematic representation of culture and drug treatments of BMDMs isolated from either young or aged mice bone marrows. **(F)** mRNA quantification of hedgehog pathway genes. **(G-H)** Representative CD80 and CD206 mRNA expression with different treatment regimens from A. (n=4). Two way ANOVA was applied followed by Tukey's test. All data are mean ± SEM (n=5): *p<0.05, **p<0.01, ***p<0.001, ****p<0.0001.

**SUPPLEMENTARY FIGURE 6**

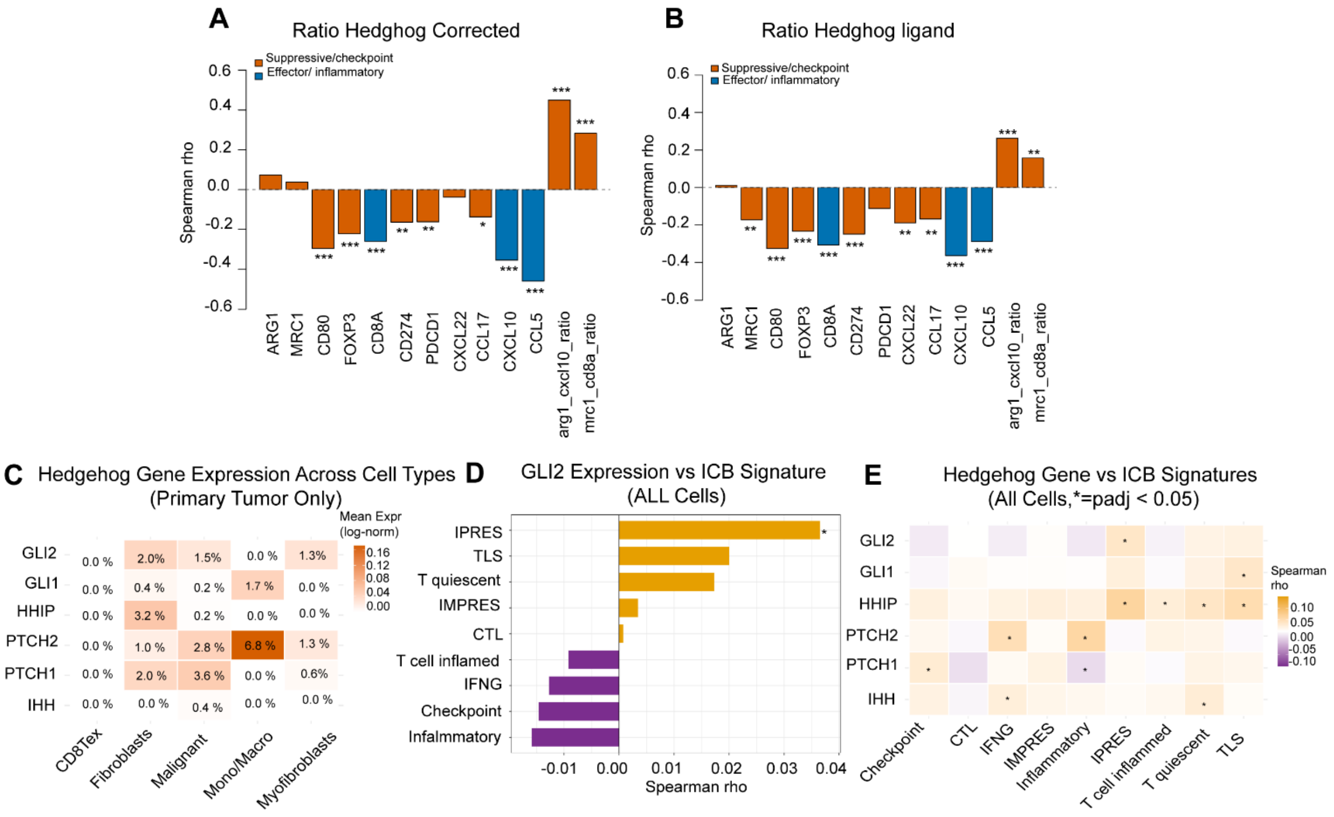


**Supplementary Figure 6.(A-B)**: Bar plots showing Spearman correlation coefficients between the (A) hedgehog pathway activation score or (B) hedgehog ligand score and selected immune-related features across TCGA ovarian tumor samples. Features include individual suppressive/checkpoint-associated genes, effector/inflammatory genes, and the composite ratios ARG1/CXCL10 and MRC1/CD8A. Orange bars indicate suppressive/checkpoint-associated features, and blue bars indicate effector/inflammatory features. (*FDR < 0.05, **FDR < 0.01, ***FDR < 0.001). **(C)** Heatmap of mean log-normalized expression of six HH pathway genes across five cell types in primary tumor tissue. Numbers within tiles indicate the percentage of cells expressing each gene. The only detected ligand, *IHH*, was restricted to malignant cells at low frequency (0.4%). *SHH* and *DHH* were not detected in this dataset (GSE130000). **(D)** Bar plot of Spearman correlation coefficients between *GLI2* expression and nine ICB response signatures across all cells (n = 11,215, GSE130000). **(E)** Heatmap of Spearman correlation coefficients between individual HH gene expression and nine ICB signatures across all cells.( GSE13000, *, $p_{adj} < 0.05$).
